# Distribution of gene tree histories under the coalescent model with gene flow

**DOI:** 10.1101/023937

**Authors:** Yuan Tian, Laura S. Kubatko

## Abstract

We propose a coalescent model for three species that allows gene flow between both pairs of sister populations. The model is designed to analyze multilocus genomic sequence alignments, with one sequence sampled from each of the three species. The model is formulated using a Markov chain representation, which allows use of matrix exponentiation to compute analytical expressions for the probability density of gene tree genealogies. The gene tree history distribution as well as the gene tree topology distribution under this coalescent model with gene flow are then calculated via numerical integration. We analyze the model to compare the distributions of gene tree topologies and gene tree histories for species trees with differing effective population sizes and gene flow rates. Our results suggest conditions under which the species tree and associated parameters are not identifiable from the gene tree topology distribution when gene flow is present, but indicate that the gene tree history distribution may identify the species tree and associated parameters. Thus, the gene tree history distribution can be used to infer parameters such as the ancestral effective population sizes and the rates of gene flow in a maximum likelihood (ML) framework. We conduct computer simulations to evaluate the performance of our method in estimating these parameters, and we apply our method to an Afrotropical mosquito data set (Fontaine et al., 2015) to demonstrate the usefulness of our method for the analysis of empirical data.

---

In multi-locus phylogenetic studies, many different evolutionary factors can cause incongruence between a gene tree and the species tree for the same set of species (Maddison, 1997). Incomplete lineage sorting (also called deep coalescence) has long been recognized to be one of the major causes of variation in gene trees across a genome (Pamilo and Nei, 1988; Takahata, 1989). Another important factor leading to discord between gene trees and the species tree is gene flow between populations following speciation (Maddison, 1997; Leaché et al., 2013; Degnan et al., 2012). With some exceptions (noted below), these two processes have been studied in isolation. When carrying out phylogenetic analyses for species that are substantially divergent, ignoring gene flow following speciation may not bias the resulting estimates. However, with the advent of large-scale genomic data sets that allow study of evolutionary relationships among closely related populations or species, the necessity of simultaneously examining these factors is becoming increasingly apparent (Eckert and Carstens, 2008; Leaché et al., 2013; Huang et al., 2014). In particular, since gene flow may easily occur between sister taxa following speciation, even in the presence of incomplete lineage sorting (Yu et al., 2011), it is necessary to incorporate these processes simultaneously into models used to analyze data for closely related species or populations.

Degnan and Salter (2005) derived the probability distribution of gene trees under the coalescent model in the absence of gene flow, and provided a method for computing this distribution that was implemented in their software, COAL. Wu (2012) provided a method of computation that was more efficient than the method of Degnan and Salter, and used this method to develop software for species tree estimation called STELLS. Although both methods model the possibility of incomplete lineage sorting using the coalescent without gene flow, the difference between the two computational approaches is in the method of enumerating possible scenarios that are consistent with a given gene tree under the model. Degnan and Salter’s approach used the concept of gene tree *histories*, which can be defined to be gene tree topologies together with an assignment of coalescent events on the gene tree topology to specific intervals of the species tree. In contrast, Wu used *ancestral configurations* to carry out the computations, where an ancestral configuration can loosely be defined as an assignment of possible states of all lineages at nodes of the species tree (see Wu (2012) for details).

The ability to compute gene tree probability distributions provided several important insights into the problem of multi-locus species tree estimation. Important among these was the realization that the gene tree topology with the highest probability need not match the species tree, a phenomenon that has led such gene trees to be called *anomalous gene trees* (Degnan and Salter, 2005; Degnan and Rosenberg, 2006). More broadly, these studies led to the realization that the incomplete lineage sorting process could result in substantial variation in the evolutionary trees for individual genes, suggesting the importance of accounting for this process in inferring species-level phylogenies. Another important insight was that the gene tree topology probability distribution identifies both the species tree topology and the speciation times (Allman et al., 2011a), which implies that if this distribution were known exactly then the species tree that produced it would also be known. This has led to the development of a collection of methods for inferring species trees from estimated gene trees (Than and Nakhleh, 2009; Liu et al., 2010; Fan and Kubatko, 2011; Wu, 2012; Mirarab et al., 2014; Bayzid et al., 2015).

Some models that incorporate both gene flow and incomplete lineage sorting jointly have also been proposed. For example, a model with incomplete lineage sorting and gene flow leading to hybrid speciation was introduced to estimate the relative parental contributions to the hybrid taxon (Meng and Kubatko, 2009) and to detect hybridization within the framework of the coalescent model (Kubatko, 2009; Gerard et al., 2011). Yu et al. (2012, 2013) proposed a model that establishes a phylogenetic network to compute the probability of gene tree topologies (Yu et al., 2012, 2013), with “horizontal” branches in the network representing gene flow or hybridization events.

Isolation-with-migration (IM) models (Hey and Nielsen, 2004; Hey, 2010) have also been used to model both population splitting and gene flow. Zhu and Yang (2012) recently used this basic model to characterize the genealogical process with both coalescence and migration. In particular, Zhu and Yang (2012) calculated the distribution of gene tree histories under an IM model with two closely related species subject to gene flow and an outgroup species, and used this probability distribution to analyze sequence data for three taxa. They used the model to obtain estimates of relevant parameters and to develop a hypothesis test for gene flow in a maximum likelihood framework. Andersen et al. (2014) used the two-population IM model with an arbitrary number of lineages in each population, and derived gene tree probability distributions under this model. They also developed procedures for inferring model parameters from sequence data in this setting.

Here we propose a new IM model for three species (not including an outgroup species) that allows gene flow between both sister populations. We formulate our model using the Markov chain representation of Hobolth et al. (2011), which allows use of matrix exponentiation to compute analytical expressions for the probability density of gene tree genealogies. We then use numerical integration to calculate the gene tree history distribution as well as the gene tree topology distribution under this coalescent model with gene flow. Our results suggest that, in contrast to the situation in the absence of gene flow, the species tree is not identifiable from the gene tree topology distribution when gene flow is present. However, the gene tree history distribution does identify the species tree topology. We also find that the gene tree history distribution can be used to infer the model parameters (such as the ancestral effective population sizes and the rates of gene flow) in a maximum likelihood (ML) framework. We conduct computer simulations to evaluate the performance of our method in estimating the model parameters. An application of our method to an Afrotropical mosquito data set (provided by Fontaine et al., 2015) is used to demonstrate the usefulness of our method for the analysis of empirical data.

## Methods

### The IM model for three species with gene flow between both sister taxa

Our proposed model for three species is shown in Figure 1. Here the three species are labeled as A, B, and C, with the species phylogeny ((A, B), C). The two ancestral species are denoted AB and ABC. The time since speciation occurred between A and B is denoted *τ*_1_, and the time since the speciation event between AB and C is denoted *τ*_2_, where *τ*_1_ and *τ*_2_ are measured by the expected number of mutations per site. The genetic data that we will analyze contain multiple loci. For every locus, we assume that one sequence was sampled from each of the three species. It is assumed that there is no recombination within a locus, and free recombination among loci. There are three possibilities for the gene tree topology relating the three sampled sequences at a locus: ((A, B), C), ((B, C), A), and ((A, C), B). Probability distributions relating to the gene tree history can be derived using Markov chains based on the structured coalescent process, as in Hobolth et al. (2011). We give the details of this approach below.

**Figure 1:**
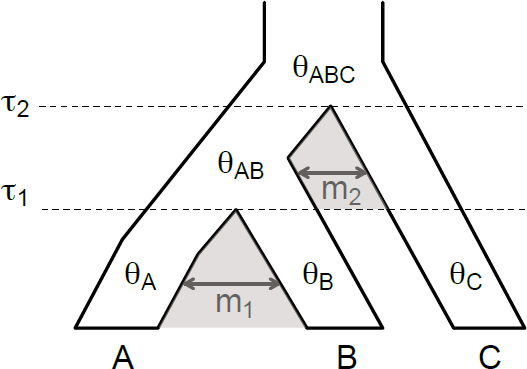
The model species tree ((A, B), C) for three species with gene flow between sister species. The speciation times are denoted as *τ*_1_ and *τ*_2_, respectively. *θ*_*A*_, *θ*_*B*_, *θ*_*C*_, *θ*_*AB*_, and *θ*_*ABC*_ are the coalescent rates for each species or ancient species. The rates of gene flow between each sister species are assumed to be equal in both directions. The gene flow rate between species A and B is *m*_1_, and the gene flow rate between species C and the ancient species AB is *m*_2_.

In our model, we consider gene flow between sister species A and B, and between sister species AB and C. More specifically, for species A and B, gene flow can occur from the present to time *τ*_1_; for species AB and C, gene flow can occur between times *τ*_1_ and *τ*_2_ (assume that there is no gene flow between species B and C, or between species A and C, after time *τ*_1_). Additionally, to simplify the calculations, we assume that the gene flow rate between sister species is the same in both directions. The parameters involved in the model are: *θ*_*A*_, *θ_B_, θ_C_, θ_AB_, θ_ABC_, m*_1_, *m*_2_, *τ*_1_, and *τ*_2_ (see Figure 1). Here *θ*_*A*_ = 4*N*_*A*_*μ*, *θ*_*B*_ = 4*N*_*B*_*μ*, *θ*_*C*_ = 4*N*_*C*_*μ*, *θ*_*AB*_ = 4*N*_*AB*_*μ*, *θ*_*ABC*_ = 4*N*_*ABC*_*μ*, where *N*_*x*_ refers to the effective population size in species x, and *μ* is the mutation rate per site. The parameters *m*_1_ and *m*_2_ are defined to be the gene flow rates between the sister species (Hobolth et al., 2011).

We use the term *gene tree genealogy* to include information for both the gene tree topology and the associated coalescent times (Degnan and Rosenberg, 2009). For a given species tree with known speciation times, we can classify gene tree genealogies into *gene tree histories* based on where coalescent events occur in relation to speciation times. We note that there are infinitely many gene tree genealogies for any number of taxa because the coalescent times associated with the genealogy are continuous parameters. However, there are finitely many gene tree histories for any species tree since there are a finite number of speciation intervals into which the coalescent times can be placed. Figure 2 will help to clarify the concept of a gene tree history.

**Figure 2:**
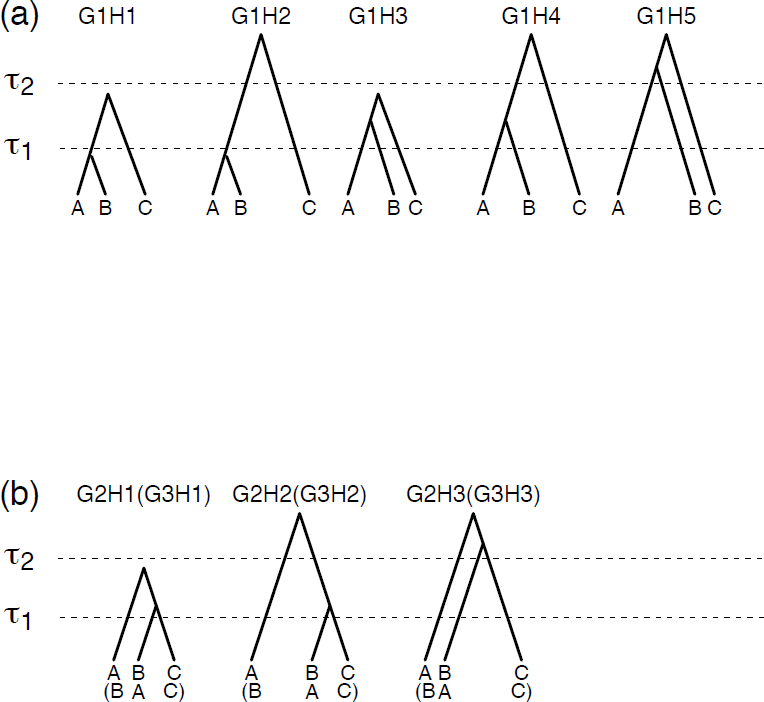
(a) Five possible gene tree histories with gene tree topology ((A, B), C) (denoted by G1). For these gene tree histories, the first (most recent) coalescent event always occurs between the lineages from species A and B. If the first coalescent event occurs before *τ*_1_, and the second coalescent event occurs between *τ*_1_ and *τ*_2_, the gene tree has history G1H1. If the first coalescent event occurs before *τ*_1_, but the second coalescent event occurs after *τ*_2_, the gene tree has history G1H2. The other three gene tree histories are denoted by G1H3, G1H4, and G1H5. (b) Three possible gene tree histories with gene tree topology ((B, C), A) (denoted by G2) and three possible gene tree histories with gene tree topology ((A, C), B) (denoted by G3, species names labeled in parentheses). For these gene tree histories, the first coalescent event occurs between the lineages from species B and C (for topology G2) or species A and C (for topology G3). If the first coalescent event occurs before *τ*_1_, and the second coalescent event occurs between *τ*_1_ and *τ*_2_, the gene tree has history G2H1 or G3H1. The other two gene tree histories are denoted as G2H2/G3H2 and G2H3/G3H3.

It is easy to see that high rates of gene flow will generate more variation in gene tree histories. Under our model in Figure 1, every species tree topology can have eleven possible histories. We denote a genealogy with gene tree topology ((A, B), C) by G1, a genealogy with gene tree ((B, C), A) by G2, and a genealogy with gene tree ((A, C), B) by G3. Within each of these there is variation in the times at which coalescent events occur and we denote the possibilities by *H*_*x*_ where × is an integer and H was chosen since these genealogies are classified to histories. As shown in Figure 2 (a), G1H1 to G1H5 show five different histories consistent with topology ((A, B), C). In Figure 2 (b), G2H1 to G2H3 show three histories consistent with topology ((B, C)), A). G3H1 to G3H3 are not shown in the figure, but they are the same as G2H1 to G2H3 with the labels for species A and B switched, leading to gene tree topology ((A, C)), B). The speciation times are labeled as *τ*_1_ and *τ*_2_ in the figure, while the coalescent time *t*_1_ (for the first nearest coalescent event from present) and *t*_2_ (for the second nearest coalescent event from present) are not labeled in the figure.

### The probability distribution of gene tree histories

Under our model, we can extend the Markov chain formulation of Hobolth et al. (2011) to calculate the probability distribution of the gene tree histories. We divide the species tree into three time periods. The first goes from the present time to *τ*_1_, and three species, A, B, and C exist during this time period; the second time period goes from *τ*_1_ to *τ*_2_, with two species AB and C; and the last goes from time *τ*_2_ to infinity, with only one species, ABC. In each time period, we can use the structured coalescent to explain the genealogical process. The method to compute the density through matrix exponentials was introduced by Hobolth et al. (2011), and the instantaneous rate matrix for two populations with gene flow was given there. We extend this method for our model, which contains three species with gene flow between two pairs of sister taxa.

#### The instantaneous rate matrix for each time period

During the first time period, from the present to *τ*_1_, gene flow can occur only between species A and B. A genealogy for a sample that includes one individual from each species has five possible states, which we denote by aac, abc, bbc, ac, bc. In our notation, aac means that two sequences are in species A, and one is in species C; abc means that one sequence is in each species; ac means that one sequence is in species A and another is in species C (here the sequences have coalesced); and so on. Note that during this time period, species C always has one lineage since there is neither gene flow nor the possibility of a coalescent event, while the ancestral populations to species A and B can experience gene flow and/or a coalescent event among the two lineages. The rates of transitions between the five states can be expressed as a 5 *×* 5 instantaneous rate matrix Q1:

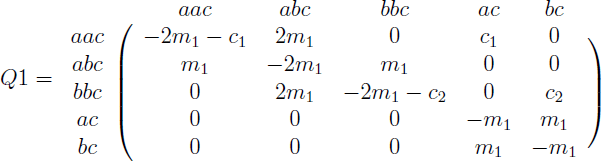

In this matrix, the coalescent parameters are defined as *c*_1_ = 2*/θ*_*A*_; *c*_2_ = 2*/θ*_*B*_; *c*_3_ = 2*/θ*_*C*_; *c*_4_ = 2*/θ*_*AB*_; *c*_5_ = 2*/θ*_*ABC*_. We use *e*^*Q*^ to denote the matrix exponential 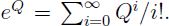. The (*j, k*)^*th*^ entry of *e*^*Q*^ is denoted as (*e*^*Q*^)_*jk*_.

Following Hobolth et al. (2011), we note that the probability density of a coalescent event at time *t*_1_ during the time period from the present time to *τ*_1_ is

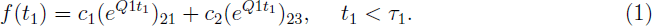

The probability density associated with the first coalescent event occurring in the ancestral population at time *t*_1_ *> τ*_1_ is

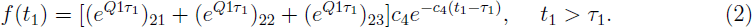

Similarly, if the first coalescent event occurs before time *τ*_1_ (*t*_1_ *< τ*_1_), a matrix Q2 can be used to compute the probability density for the second coalescent event at time *t*_2_, as follows. First denote species AB as population d, and species ABC as population e. The five possible genealogy states starting at time *τ*_1_ now become dd, dc, cc, d, c, and the associated rate matrix is

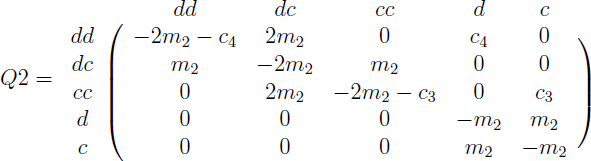

During the time period from *τ*_1_ to *τ*_2_, the probability density for the coalescent event at time *t*_2_ *< τ*_2_ is

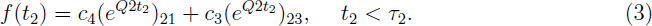

and the density for the coalescent event at time *t*_2_ occuring in the ancestral population at time *t*_2_ *> τ*_2_ is

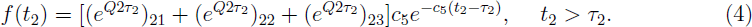

The system becomes more complicated if the first coalescent event occurs after time *τ*_1_ (*t*_1_ *> τ*_1_). In that case, three lineages exist after time *τ*_1_ and all of them can migrate between species AB and C. A 13 *×* 13 rate matrix Q3 can be built to calculate the gene tree density. We label the state of each lineage sequentially. For instance, ddc refers to the first two lineages being in population d (species AB), while the third lineage is in population c (species C). There are 13 possible states, as shown in matrix Q3,

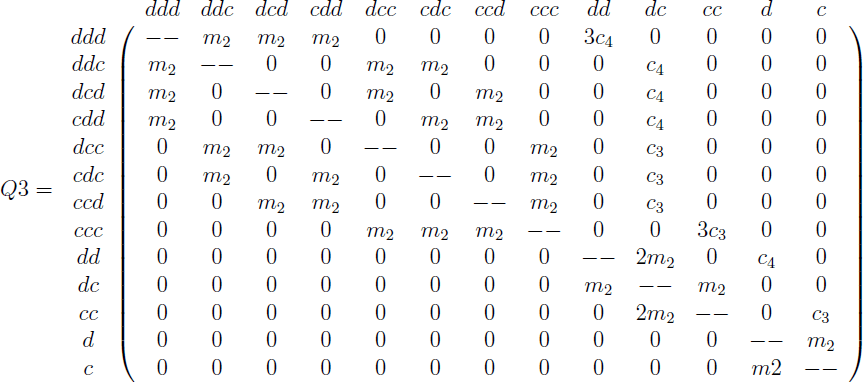

where the diagonal entries (the ‘- -’ above) are set to the negative sum of the corresponding row.

In our model, we use Q3 for the time period between time *τ*_1_ and *t*_1_ when the first coalescent event occurs after time *τ*_1_ (*t*_1_ *> τ*_1_). At the beginning of this time period, there are 3 lineages with state ddc. At the end of this time period, since gene flow can occur between populations d and c, any state with two or three lineages is possible.

#### The probability distribution of the gene tree histories

Using the results above, the probability distribution of the gene tree histories can be calculated. Recall that there are three different gene tree topologies, denoted as G1, G2, and G3 for topologies ((A, B), C), ((B, C, A), and ((A, C), B), respectively (Figure 2). Because coalescent events can occur in different intervals on the species tree, there are multiple gene tree histories that are consistent with each gene tree topology. For example, as shown in Figure 2(a), gene tree topology G1 can result from 5 different histories, labelled as G1H1, G1H2, and so on. Similarly, G2 and G3 are both consistent with 3 different histories. The probability of each history will be calculated separately using the Markov chain formulation above. We give example calculations for a few histories below. The remaining calculations are given in Appendix.

For G1H1, the first coalescent event occurs between the present and time *τ*_1_, while the second coalescent event occurs between time *τ*_1_ and *τ*_2_, thus *t*_1_ *< τ*_1_ *< t*_2_ *< τ*_2_. To derive the joint density of the coalescent times for this gene tree history, we first consider the time range between the present and time *τ*_1_. Since there is no gene flow between species C and any other species during this time, we only need to calculate the probability that the two lineages in species A and B coalesce before time *τ*_1_. Due to the possibility of gene flow between species A and B, the coalescent event can happen in either species A or B. The whole process can be described as three lineages that start in state abc, and then right before the first coalescent event, the state becomes either aac or bbc. Thus from (**??**), we can derive

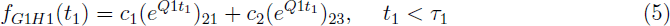

Similarly, the second coalescent event occurs between time *τ*_1_ and *τ*_2_, and the process is assumed to start from state dc. Right before the second coalescent event, the state changes to dd or cc. From the previous function (**??**), we have

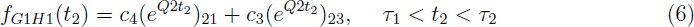

The joint distribution for both *t*_1_ and *t*_2_ is then

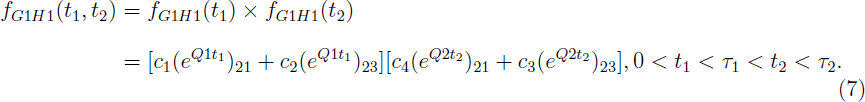

To find the marginal probability of gene tree history G1H1, we integrate out the gene tree coalescent times,

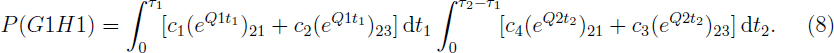

For G1H2, the first coalescent event occurs between the present and time *τ*_1_, while the second coalescent event occurs after time *τ*_2_, and thus *t*_1_ *< τ*_1_ *< τ*_2_ *< t*_2_. The only difference in the calculations for G1H2 is that no coalescent event occurs between time *τ*_1_ and *τ*_2_. The probability that the second coalescent event occurs before time *τ*_2_ is

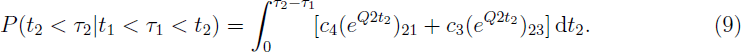

Thus, the probability that the second coalescent event occurs after time *τ*_2_ is

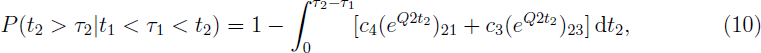

and

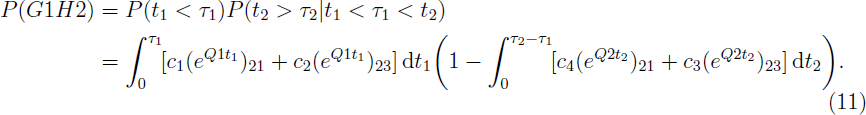

For G1H3, the first coalescent event occurs after time *τ*_1_, while the second coalescent event occurs before time *τ*_2_, and thus *τ*_1_ *< t*_1_ *< t*_2_ *< τ*_2_. From the present to time *τ*_1_, no coalescent event occurs, and the probability is

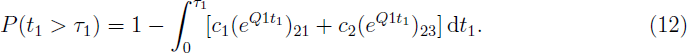

After time *τ*_1_, things are more complicated. There are four distinct ways in which the two coalescent events can happen. In all four cases, the process starts in state ddc, which means that two lineages are in population d and one is in population c. The first case denoted by G1H3C1, goes from state ddc to ddd, and then a coalescent event occurs and the state is dd. The final coalescent event can occur either from this state, or by changing to state cc. To model this, we use Q3 to calculate the change from state ddc to ddd, and use Q2 for the change from state dd to dd or cc. The joint density function is

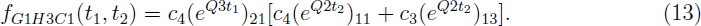

Notice that in the above density function, there is no multiplier for *c*_4_ in the first coalescent process. Although there are three possible lineage combinations that can coalesce at time *t*_1_, only one of them (the lineages that come from species A and species B) will maintain the gene tree topology ((A, B), C).

Similarly, we can write the density functions for the other three possibilities. The second possibility has the sequence of states ddc - ddc - dc - dd/cc, with corresponding density function

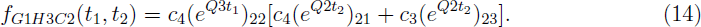

The third possibility has the sequence of states ddc - ccc - cc - dd/cc, with corresponding density function

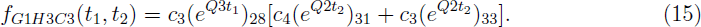

Finally, the fourth possibility has the sequence of states ddc - ddc - dc - dd/cc, with corresponding density function

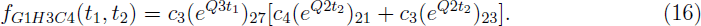

Thus, the overall density function for *t*_1_ and *t*_2_ for history G1H3 is

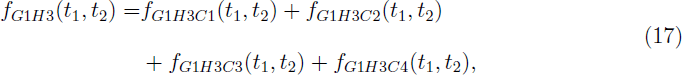

and the marginal probability of G1H3 is

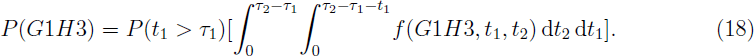

The probabilities of all other gene tree histories can be calculated similarly. We give the details in Appendix.

#### Implementation details and parameter scaling

To calculate the probability for each gene tree history requires computing integrals or double integrals (for histories G1H3, G2H1 and G3H1). To do this, we used Gaussian Quadrature for one-dimensional integration, and iterated it for two-dimensional integration. The method is implemented in a C program COALGF Calculator (COALGF) that directly calculates the probabilities for all eleven gene tree histories as well as the three gene tree topologies. The required input parameters in COALGF are the coalescent rates *c*_1_, *c*_2_, *c*_3_, and *c*_4_ (for population A, B, C, and AB, respectively); the gene flow rates *m*_1_ and *m*_2_ (*m*_1_ for the gene flow rate between population A and B, *m*_2_ for the gene flow rate between population AB and C; we assume equal rates of gene flow to and from sister species); and the speciation times *τ*_1_ and *τ*_2_.

To simplify the calculation, all parameters are scaled in COALGF so that they are proportional to a selected *c*_0_ = 2*/θ*_0_. Consider the probability of the first gene tree history in (**??**). Using matrix *Q*1 as an example, note that for *c*_0_ = 2*/θ*_0_ we can write

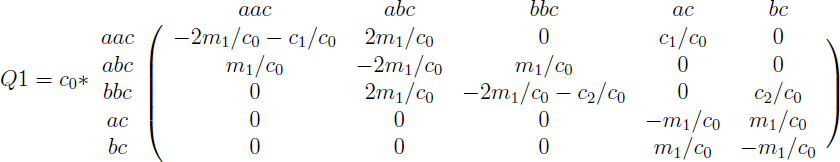

Note that *Q*1 = *c*_0_ *\* Q*1*′*, where *Q*1*′* is the new matrix above, with all coalescent rates and gene flow rates scaled by *c*_0_, and with 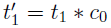. Similarly we can scale all coalescent rates by dividing *c*_0_, and all speciation times by multiplying *c*_0_. In the following results, we use *θ*_*x*_, *c*_*x*_, *m*_*x*_, and *τ*_*x*_ as the original parameters, which are not scaled by 2*/θ*_0_. For the scaled parameters, we use *C*_*x*_ = *c*_*x*_/*c*_0_, *M*_*x*_ = *m*_*x*_/*c*_0_, and *T*_*x*_ = *τ*_*x*_ * *c*_0_ as the scaled coalescent parameters, gene flow parameters, and speciation times, respectively.

### Maximum likelihood parameter estimation

Using the results of last section, the exact probabilities for the eleven gene tree histories can be calculated for any species tree with three species. Thus, given a data set consisting of observations of gene tree histories, these data can be viewed as a sample from a multinormial distribution with eleven categories and with probabilities as derived in the previous section. The likelihood of the data can thus be used to obtain maximum likelihood estimates of the coalescent parameters. We use simulation to assess the performance in estimating these parameters.

Three simulation studies were carried out using the software ms (Hudson, 2002) and seq-gen (Rambaut and Grassly, 1997). The first simulates gene trees directly, while the second and the third simulate 500bp and 1000bp DNA sequences, respectively. The DNA sequence data sets are analyzed by PAUP* (Swofford, 2003) to estimate gene trees under the maximum likelihood criterion. In each simulation study, we select a varying number of loci (ranging from 50 to 100,000) to assess our model. All data are simulated under the fixed species tree ((A, B), C), with *θ*_*A*_ = *θ*_*B*_ = *θ*_*C*_ = *θ*_*AB*_ = 0.005, *m*_1_ = *m*_2_ = 200, *τ*_1_ = 0.004, and *τ*_2_ = 0.006, which were chosen based on Zhu and Yang (2012). After scaling by *θ*_0_ = 0.005, we have: *C*_1_ = *C*_2_ = *C*_3_ = *C*_4_ = 1, *M*_1_ = *M*_2_ = 0.5, *T*_1_ = 1.6, and *T*_2_ = 2.4.

For each simulated data set with K loci, the frequency of the *x*^*th*^ gene tree history is recorded as *k*_*x*_, *x* = 1, 2, *… ,* 11. In order to estimate the model parameters, we consider two methods for searching parameter space to find the maximum likelihood estimate (MLE), both based on a grid search. The first assumes that *M*_1_ = *M*_2_ = *M* and *C*_1_ = *C*_2_ = *C*_3_ = *C*_4_ = *C*, with C varying from 0 to 2 (we used 200 equal spaced values), and M varying from 0 to 10 (we used 200 different values on the log scale). The other method assumes that *C*_1_ = *C*_2_ = *C*_3_ = *C*_4_ = 1, and varies both *M*_1_ and *M*_2_ from 0 to 10 (we used 200 different values on the log scale). Note that although we could consider *M*_1_, *M*_2_, and *C* at the same time, in the simulation study, we only considered two parameters at a time in order to reduce the computational burden and to run more replications. For the empirical data, these parameters are estimated together.

For both methods, 40,000 species trees were tested. For each species tree, the exact probability distribution of the 11 gene tree histories was calculated and the likelihood for each simulated data set was calculated (Wang and Hey, 2010; Zhu and Yang, 2012). The parameters with the highest likelihood are the maximum likelihood estimates. We simulated 1,000 replications for each simulation condition, and computed the average and the standard deviation of the MLEs of the model parameters over these replicates for each simulation condition.

### Application of the model in an empirical Afrotropical mosquito data set

Fontaine et al. (2015) reported pervasive autosomal gene introgression in several Afrotropical mosquito sibling species. In their study, the species branching order of seven Afrotropical mosquito sibling species was identified, and the times between speciation events were also estimated (Fontaine et al., 2015). We selected three of these species, *Anopheles coluzzii* (*An. col*), *Anopheles gambiae* (*An. gam*), and *Anopheles arabiensis* (*An. ara*), to serve as the species A, B, and C to test our model. We also selected an outgroup species, *Anopheles christyi* (*An. chi*), to root the gene trees. Based on Fontaine et al. (2015), the estimated species tree for the four species is (((*An. col*, *An. gam*), *An. ara*), *An. chr*). The speciation time between *An. col* and *An. gam* is 0.54 million years ago (Myr), and the speciation time between *An. ara* and the ancestor of *An. col* and *An. gam* is 1.85 Myr (Fontaine et al., 2015).

We used the whole genome alignment of the reference assemblies from the members of the *Anopheles gambiae* species complex (Fontaine et al., 2015), and selected data from chromosome 2L to analyze. In total, 24,921 gene trees were constructed from 1 kb non-overlapping windows across the alignments by PAUP* using maximum likelihood. Based on the speciation times given above, the frequencies of the gene tree histories were recorded. Assuming that all three selected *Anopheles gambiae* species have equal effective population sizes, the effective population size, the gene flow rate between *An. col* and *An. gam*, and the gene flow rate between *An. ara* and the ancestor of *An. col* and *An. gam* were estimated using maximum likelihood.

## Results

### Gene flow between sister species produces different distributions of gene tree histories

We calculated the probability distribution of gene tree histories using the method described above under a set of species trees with different parameter values (Figure 3). In the figure, the different gene tree histories are indicated with different colors, and the vertical height of each colored bar shows the probability of that history. Histories are grouped according to their topology, and since the probability of gene tree topologies G2 and G3 are always equal, only one bar is shown in the figure (labeled G2/3). It is clear that the sum of the probability of G1 and twice of the probability of G2/3 is equal to 1 under each species tree setting. The effect of gene flow on the distribution of gene tree histories is explored under two different conditions: all current and ancestral populations have equal effective population sizes (Figure 3(a)), and current and ancestral populations have unequal effective population sizes (Figure 3(b)).

**Figure 3:**
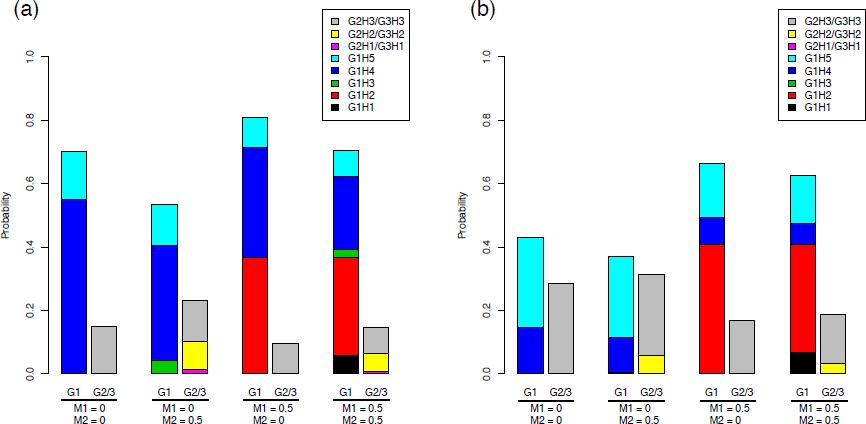
The probability distribution of the gene tree histories under species trees with scaled speciation times *T*_1_ = 1.6, *T*_2_ = 2.4 (*τ*_1_ = 0.004, *τ*_2_ = 0.006). Each gene tree history is denoted by a different color as shown in the figure. The probability of topology G1 is shown by the height of the column labeled G1; the height of the column labeled G2/3 shows the equal probability of the topologies G2 and G3. Thus, *P*(*G*1) + 2*P*(*G*2*/*3) = 1. The two sets of scaled coalescent rates are *C*_1_ = *C*_2_ = *C*_3_ = *C*_4_ = 1 (scaled by *θ*_0_ = 0.005) in panel (a), and *C*_1_ = 1, *C*_2_ = *C*_3_ = 0.5, *C*_4_ = 0.2 (scaled by *θ*_0_ = 0.005) in panel (b). Each panel contains four cases of different gene flow rates: 1, no gene flow (*M*_1_ = *M*_2_ = 0); 2, no gene flow between species A and B (*M*_1_ = 0, *M*_2_ = 0.5); 3, no gene flow between species C and the ancient species AB (*M*_1_ = 0.5, *M*_2_ = 0); 4, equal rates of gene flow in both sister species (*M*_1_ = *M*_2_ = 0.5).

In Figure 3 (a), the effective population sizes of species A, B, C, and AB (the ancestor of species A and B) are assumed to be equal. Assuming that *θ*_0_ = 0.005, *τ*_1_ = 0.004, and *τ*_2_ = 0.006 (Zhu and Yang, 2012), the coalescent parameters were scaled to *C*_1_ = *C*_2_ = *C*_3_ = *C*_4_ = 1, and the relative speciation times were scaled to *T*_1_ = 1.6, and *T*_2_ = 2.4. When there is no gene flow between either pair of sister species (Figure 3 (a) *M*_1_ = 0, *M*_2_ = 0), only three histories are possible (G1H4, G1H5, and G2H3/G3H3; see Figure 2), since the first coalescent event cannot occur before speciation time *T*_1_, and the second coalescent event cannot occur before speciation time *T*_2_. When gene flow can occur between species AB and C, the second coalescent event could also occur between speciation time *T*_1_ and *T*_2_, and thus G1H3, G2H1/G3H1 and G2H2/G3H2 (see Figure 2) will also have positive probability. In that case, topology G1 contains three different histories, while topologies G2/G3 also have three different histories (Figure 3 (a) *M*_1_ = 0, *M*_2_ = 0.5). However, if there is no gene flow between species AB and C, but gene flow is possible between species A and B (Figure 3 (a) *M*_1_ = 0.5, *M*_2_ = 0), again only one history (G2H3/G3H3) is possible for topologies G2/3, since the second coalescent event cannot occur before speciation time *T*_2_, but the first coalescent event can occur before speciation time *T*_1_. The gene tree topology G1 will still contain three histories, but these are different than the three that appear with *M*_1_ = 0, *M*_2_ = 0.5. When gene flow is possible between both pairs of sister species (species A & B and species AB & C), all of the histories in Figure 2 have positive probability (Figure 3 (a) *M*_1_ = 0.5, *M*_2_ = 0.5).

While the gene tree history distribution shows a clear pattern when all effective population sizes are assumed to be equal (Figure 3 (a)), the distribution changes when these parameters differ across the species tree. In Figure 3 (b), we used a set of species trees with scaled coalescent parameters *C*_1_ = 1, *C*_2_ = *C*_3_ = 0.5, *C*_4_ = 0.2 along with all of the other parameter combinations in Figure 3 (a). The ratio of the effective population sizes considered here was selected based on Burgess and Yang’s (2008) paper in which hominoid ancestral population sizes were estimated. With regard to which settings lead to positive probabilities associated with various gene tree histories, the patterns in Figure 3 (a) and (b) are generally consistent. Except for the case in which *M*_2_ ≠ 0, histories G1H3 and G2H1/G3H1 have extremely small probabilities (Figure 3 (a) (b) *M*_1_ = 0, *M*_2_ = 0.5; *M*_1_ = 0.5, *M*_2_ = 0.5). This result is not surprising, because the length of time between the two speciation times *T*_1_ and *T*_2_ is relatively small (*T*_2_ − *T*_1_ = 0.8). Histories G1H3 and G2H1/G3H1 only arise when both coalescent events occur between *T*_1_ and *T*_2_ (Figure 2). With a small time span between *T*_1_ and *T*_2_ as well as unequal effective population sizes, these probabilities become very small. If the time span between *T*_1_ and *T*_2_ is longer, as in Figure S1 (b), *T*_1_ = 2 and *T*_2_ = 4, it is clear that all five histories show up in topology G1, and all three histories appear in G2/3, when *M*_2_ *≠* 0. Note that all other parameters (coalescent rates and gene flow rates) are the same in Figure 3 (a) and Figure S1 (b). Finally, through we observe a similar pattern for the gene tree history probabilities for the species trees with equal or unequal population sizes, the magnitude of the probability of each history varies a lot. For example, in Figure 3 (a), G1H4 is one of the dominant histories, while in Figure 3 (b), the probability of G1H4 is much smaller and the probabilities of G1H5 and G2H3/G3H3 are dominant.

We also considered another set of speciation times, *T*_1_ = 2 and *T*_2_ = 4, which has a much longer time span between the two speciation events (Figure S1). Comparing Figure 3(a) and Figure S1(a) shows that with longer speciation times, the probabilities of histories G1H5 and G2H3/G3H3 clearly decrease, because the increase in speciation times decreases the probability that both coalescent events occur before the earliest speciation time. Interestingly, if there is no gene flow between either pair of sister species, the longer speciation times will greatly decrease the probability of observing the “incorrect” gene tree topology (G2 or G3). Gene flow between only species A and B, but not species AB and C, will lead to a similar distribution because the topology G2/G3 is still composed of just one gene tree history (G2H3/G3H3) and the probability of this history decreases with longer speciation times. However, if ancient gene flow exists between species AB and C, the distribution of gene tree topologies does not change a lot, but the distribution of gene tree histories shows some clear changes (compare Figure 3 (a) (b) and Figure S1 (a) (b)).

We also considered a second level for the rate of gene flow. In Figure S1(c)-(f), the rate of gene flow was set to 2 when it was present (in contrast to the rate of 0.5 used in Figure 3 and Figure S1(a), (b)). For the same effective population sizes and the same speciation times, we find that changing the rate of gene flow from 0.5 to 2 does not have a huge effect on the distribution of gene tree histories. More extreme values of the rate of gene flow will be discussed in the following sections.

### Variation in the gene tree history distribution as a function of the rate of gene flow

We considered the change in the probabilities of individual histories as a function of the gene flow rate when all other parameters were held constant. In each subplot of Figure 4 and Figures S2 - S5, *M*_1_ is held constant (at four different levels *M*_1_ = 0.001; 0.5; 2; 20, corresponding to the rows), while the value of *M*_2_ changes from 0 to 1, 000. The effective population sizes and speciation times both have two different levels. For the effective population sizes, one setting is that all effective population sizes are equal, thus the coalescent rates were set to *C*_1_ = *C*_2_ = *C*_3_ = *C*_4_ = 1 (Figure 4 (a) - (i), Figures S2, S3), and the other setting is a species tree with unequal effective population sizes, *C*_1_ = 1, *C*_2_ = *C*_3_ = 0.5, *C*_4_ = 0.2 (Figure 4 (j) - (l), Figures S4, S5). For the speciation times, a set of relatively shorter speciation times [*T*_1_ = 1.6; *T*_2_ = 2.4 (Figure 4 (a) - (f) (j) - (l), Figures S2, S4)] and a set of longer speciation times [*T*_1_ = 2; *T*_2_ = 4 (Figure 4 (g) - (i), Figures S3, S5)] are used. Figure 5 and Figures S6 - S9 have similar parameter settings, but in these figures, *M*_2_ is held constant at four different levels, while the value of *M*_1_ is changed from 0 to 1, 000. In all of these figures that plot the distribution of gene tree histories against the rate of gene flow, the first column shows the distribution of the five gene tree histories with topology G1=((A,B),C), the second column shows the distribution of the three gene tree histories with the other two topologies G2/G3, and the last column shows the distribution of the three possible gene tree topologies (G1, G2, and G3). Note that in the last column, the probabilities of the topologies G2 and G3 are always equal.

**Figure 4:**
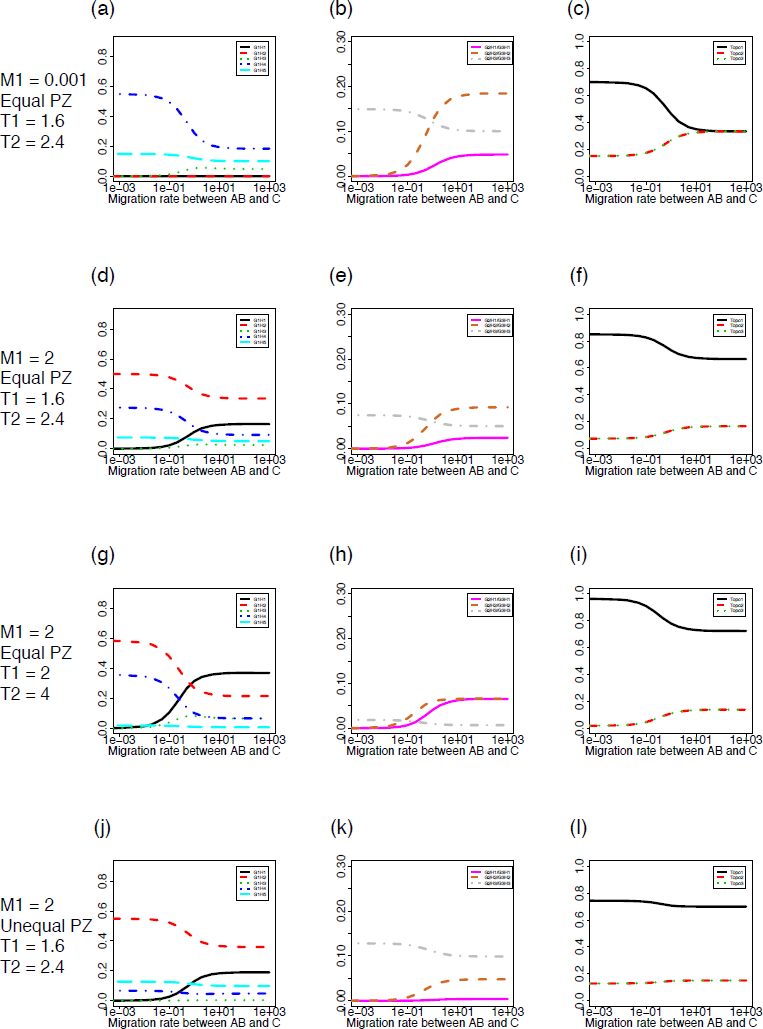
Probability distributions of the gene tree histories for the model of three species with gene flow between sister species. The probabilities of each gene tree history (y-axis) were plotted against the gene flow rate between species C and the ancient species AB, *M*_2_ (x-axis; shown on a log scale). Panels (a), (d), (g), and (j) show the probabilities of the five gene tree histories (G1H1 - G1H5) with topology G1. Panels (b), (e), (h), and (k) show the probabilities of the three gene tree histories (G2H1/G3H1 - G2H3/G3H3) with topology G2 or G3. Panels (c), (f), (i), and (l) show the probabilities of the three gene tree topologies (G1, G2 and G3) with *P*(*G*2) = *P*(*G*3), and *P*(*G*1) + *P*(*G*2) + *P*(*G*3) = 1. The four sets of parameter values are *C*_1_ = *C*_2_ = *C*_3_ = *C*_4_ = 1, *T*_1_ = 1.6, *T*_2_ = 2.4, *θ*_0_ = 0.005, *M*_1_ = 0.001 for panels (a) - (c); *C*_1_ = *C*_2_ = *C*_3_ = *C*_4_ = 1, *T*_1_ = 1.6, *T*_2_ = 2.4, *θ*_0_ = 0.005, *M*_1_ = 2 for panels (d) - (f); *C*_1_ = *C*_2_ = *C*_3_ = *C*_4_ = 1, *T*_1_ = 2, *T*_2_ = 4, *θ*_0_ = 0.01, *M*_1_ = 2 for panels (g) - (i); and *C*_1_ = 1, *C*_2_ = *C*_3_ = 0.5, *C*_4_ = 0.2, *T*_1_ = 1.6, *T*_2_ = 2.4, *θ*_0_ = 0.005, *M*_1_ = 2 for panels (j) - (l).

**Figure 5:**
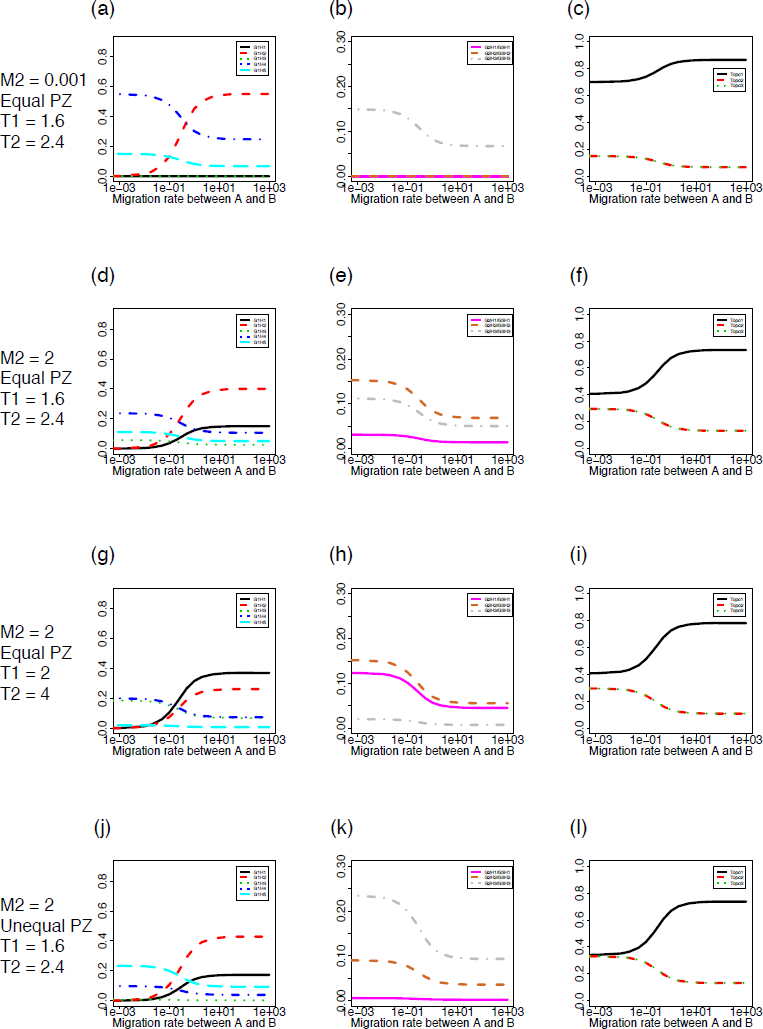
Probability distributions of the gene tree histories for the model of three species with gene flow between sister species. The probabilities of each gene tree history (y-axis) were plotted against the gene flow rate between species A and species B, *M*_1_ (x-axis; shown on a log scale). The four sets of parameter values are *C*_1_ = *C*_2_ = *C*_3_ = *C*_4_ = 1, *T*_1_ = 1.6, *T*_2_ = 2.4, *θ*_0_ = 0.005, *M*_2_ = 0.001 for panels (a) - (c); *C*_1_ = *C*_2_ = *C*_3_ = *C*_4_ = 1, *T*_1_ = 1.6, *T*_2_ = 2.4, *θ*_0_ = 0.005, *M*_2_ = 2 for panels (d) - (f); *C*_1_ = *C*_2_ = *C*_3_ = *C*_4_ = 1, *T*_1_ = 2, *T*_2_ = 4, *θ*_0_ = 0.01, *M*_2_ = 2 for panels (g) - (i); and *C*_1_ = 1, *C*_2_ = *C*_3_ = 0.5, *C*_4_ = 0.2, *T*_1_ = 1.6, *T*_2_ = 2.4, *θ*_0_ = 0.005, *M*_2_ = 2 for panels (j) - (l).

When there is no gene flow between species A and B (i.e., *M*_1_ = 0.001; Figure 4 (a) - (c)) or between species AB and C (i.e., *M*_2_ = 0.001;Figure 5 (a) - (c)), different gene tree histories will have positive probability compared with the case in which gene flow is present. For example, comparing Figure 4 (a) - (c) and (d) - (f), we see that some gene tree histories have positive probability only when there is gene flow (G1H1 and G1H2). Comparing Figure 5 (a) - (c) and (d) - (f), we find the same pattern, but with different gene tree histories (G1H1 and G1H3). Notably, determining whether or not gene flow occurred based on the frequencies of gene tree topologies would be difficult, especially when the rate of gene flow is not large (Figure 4 (c), (f); Figure 5 (c), (f)). Another interesting fact is that if the rate of gene flow is held constant for one pair of sister species, the gene tree history distribution can change completely as the other gene flow rate changes from a small value to a large value (Figure 4 and Figure 5). These results suggest that the distribution of gene tree histories depends highly on the magnitude of gene flow.

In addition to gene flow, two other factors may affect the distribution of gene tree histories. The first factor is the speciation time. Figure 4 (d) - (f) and (g) - (i) show the differences in the gene tree history distributions for relatively smaller speciation times (*T*_1_ = 1.6; *T*_2_ = 2.4) and for larger speciation times (*T*_1_ = 2; *T*_2_ = 4). When *M*_2_ is low (less than 0.1), the distributions of gene tree histories under different speciation times show a very similar pattern with slightly different values. However, when *M*_2_ is larger, the distributions of gene tree histories with the two sets of different speciation times have very different patterns. It is quite interesting to notice that the topology distribution differs more when *M*_2_ is small (less than 0.1), but becomes more similar as *M*_2_ becomes larger (Figure 4 (f) and (i)). This effect is even larger in Figure 5 (d) - (f) and (g) - (i), when *M*_2_ is held constant 25 and *M*_1_ varies.

The other factor that may affect the distribution of gene tree histories is the effective population size, which determines the rate at which coalescent events occur. We observe that, for the two different sets of coalescent rates we considered (equal effective population sizes [*C*_1_ = *C*_2_ = *C*_3_ = *C*_4_ = 1] and unequal effective population sizes [*C*_1_ = 1, *C*_2_ = *C*_3_ = 0.5, *C*_4_ = 0.2]), the gene tree topology distribution is affected less by the change of effective population sizes than the gene tree history distribution (Figure 4 (d) - (f) and (j) - (l); Figure 5 (d) - (f) and (j) - (l)). Complete comparisons for different rates of gene flow, speciation times, and effective population sizes are shown in Figures S2 - S9.

### Different distributions of gene tree histories may share an identical gene tree topology distribution

Under the typical coalescent model without the possibility of gene flow following speciation, the distribution of gene tree topologies can be used to estimate the species tree topology and branch lengths (Allman et al., 2011b). However, in the presence of gene flow, the gene tree topology probabilities change, and we might ask whether this distribution alone is sufficient to identify whether or not gene flow has occurred. Our overall finding is that many different distributions of gene tree histories arising from different species trees may share an identical gene tree topology distribution, which indicates that the information about gene tree topology probabilities alone is not enough to estimate species trees in the presence of gene flow.

For example, Figure 6 shows eight species trees (*S_*1 – *S_*8) that all have the same set of scaled speciation times, *T*_1_ = 1.6 and *T*_2_ = 2.4. Trees *S_*1 – *S_*4 have equal effective population sizes with scaled coalescent parameters *C*_1_ = *C*_2_ = *C*_3_ = *C*_4_ = 1, while species trees *S_*5 – *S_*8 have unequal effective population sizes with scaled coalescent parameters *C*_1_ = 1, *C*_2_ = *C*_3_ = 0.5, *C*_4_ = 0.2. The eight species trees differ in the rates of gene flow, *M*_1_ and *M*_2_. For instance, *S_*1 shows the case of no gene flow (*M*_1_ = *M*_2_ = 0.001), while *S_*5 shows the case that gene flow occurs only between species A and B (*M*_1_ = 0.798; *M*_2_ = 0.001). Different levels of gene flow are also reflected in this figure: *S_*2 has small gene flow rates (*M*_1_ = 0.5; *M*_2_ = 0.544); *S_*8 has medium gene flow rates (*M*_1_ = 2; *M*_2_ = 5); and *C* 4 has fairly large gene flow rates (*M*_1_ = 20; *M*_2_ = 28). As labeled in the figure, different colors show the probabilities of different gene tree histories. For each species tree, the probabilities of the eleven gene tree histories sum up to 1. The two solid black vertical lines in Figure 6 divide the total probability into the three probabilities corresponding to the gene tree topologies (from left to right, G1, G2, and G3), which are *identical* for all eight cases. Under this identical topology distribution (*P*(*G*1) = 0.7, *P*(*G*2) = *P*(*G*3) = 0.15), the distributions of gene tree histories are very different for these eight species trees.

**Figure 6:**
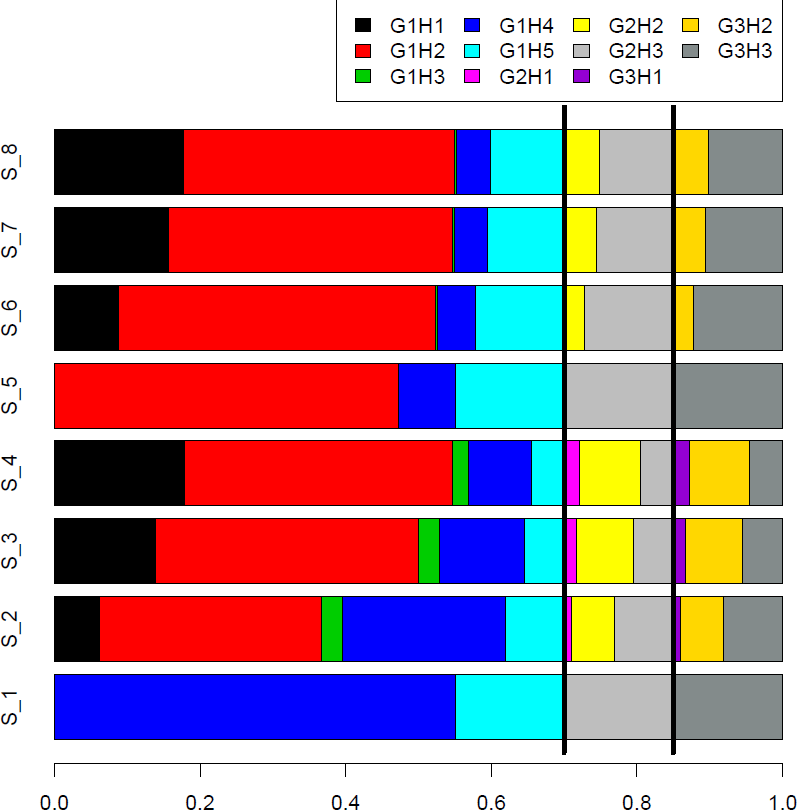
Gene tree history distributions for eight species trees (rows labeled *S_*1 – *S_*8) with different coalescent rates and gene flow rates. The x-axis is the probability associated with individual histories (denoted by colors), and the total probability for each case sums up to 1. The two solid black lines indicate the probability of each of the three gene tree topologies, from left to right *P* (*G*1) = 0.7, and *P* (*G*2) = *P* (*G*3) = 0.15. All eight species trees have scaled speciation times *T*_1_ = 1.6, *T*_2_ = 2.4 (scaled by *θ*_0_ = 0.005). The two sets of coalescent rates are *C*_1_ = *C*_2_ = *C*_3_ = *C*_4_ = 1 for species trees *S_*1 − *S_*4, and *C*_1_ = 1, *C*_2_ = *C*_3_ = 0.5, *C*_4_ = 0.2 for species trees *S_*5 − *S_*8. All species trees have different rates of gene flow: *S_*1: *M*_1_ = *M*_2_ = 0; *S_*2: *M*_1_ = 0.5, *M*_2_ = 0.544; *S_*3: *M*_1_ = 2, *M*_2_ = 2.23; *S_*4: *M*_1_ = 20, *M*_2_ = 28; *S_*5: *M*_1_ = 0.796, *M*_2_ = 0; *S_*6: *M*_1_ = 1.337, *M*_2_ = 0.5; *S_*7: *M*_1_ = 1.894, *M*_2_ = 2; *S_*8: *M*_1_ = 2, *M*_2_ = 5.

Notably, these eight species trees are not the only species trees that share this particular gene tree topology distribution, and this gene tree topology distribution is not the only one which can be generated by multiple species trees. However, despite the implied non-identifiability of gene flow based only on the topology distribution, we note that the gene tree history distributions appear to be distinct in these cases. Because of this, the information provided by the distribution of gene tree histories can be used to estimate the parameters (effective population sizes and gene flow rates) in the coalescent model with gene flow, as we show in the following two sections.

### Simulation studies for maximum likelihood parameter estimation

To assess the performance of our model in estimating the rates of coalescence and gene flow, we carried out three simulation studies. In the first simulation study, we simulated gene trees directly with a varying number of loci ranging from 50 to 100,000. Gene trees were simulated under the fixed species tree ((A, B), C), with *θ*_*A*_ = *θ*_*B*_ = *θ*_*C*_ = *θ*_*AB*_ = 0.005, *m*_1_ = *m*_2_ = 200, *τ*_1_ = 0.004, and *τ*_2_ = 0.006, corresponding to scaled parameters *C*_1_ = *C*_2_ = *C*_3_ = *C*_4_ = 1, *M*_1_ = *M*_2_ = 0.5, *T*_1_ = 1.6, and *T*_2_ = 2.4. Assuming that *M*_1_ = *M*_2_ = *M* and *C*_1_ = *C*_2_ = *C*_3_ = *C*_4_ = *C*, we evaluated the likelihood of 40,000 species trees (each differing in the values of *M* and *C* but with the topology and branch lengths fixed) to find the MLEs of *M* and *C*. Figure 7 shows the results for one simulated data set in this simulation study for which 1000 gene trees were simulated with *C* = 1 and *M* = 0.5. It is clear that the MLEs *Ĉ* = 0.99 and 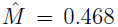 = 0.468, marked with a white ’X’ in Figure 7, are close to the true values.

**Figure 7:**
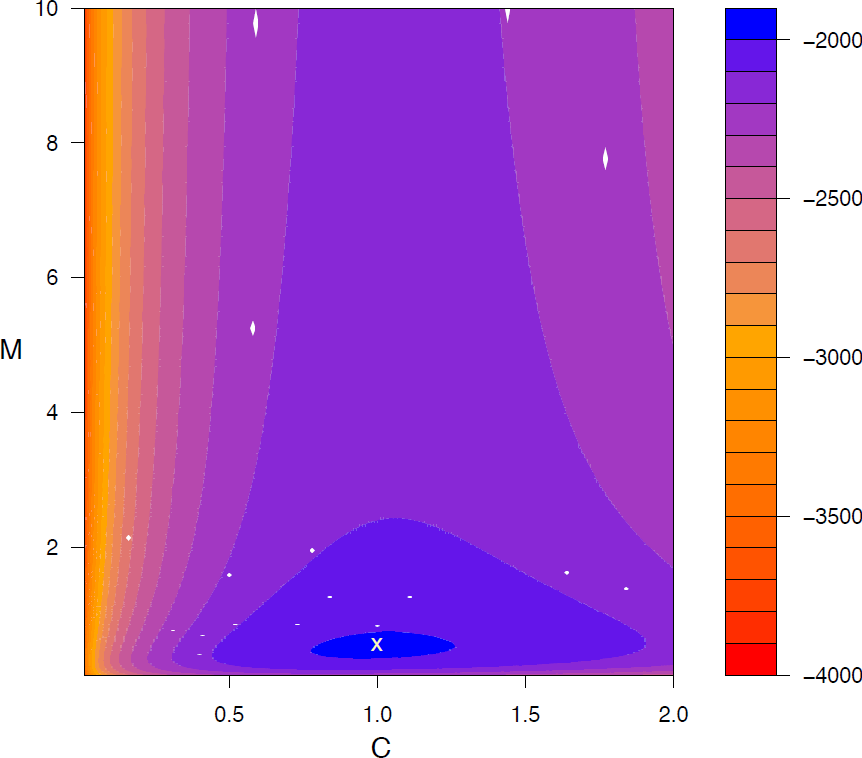
Contour plot for one simulated data set showing the log likelihood as a function of the scaled coalescent rate C and the gene flow rate M, assuming that *C*_1_ = *C*_2_ = *C*_3_ = *C*_4_ = *C* and *M*_1_ = *M*_2_ = *M*. DNA sequence data were simulated for 1000 loci for species tree ((A, B), C) with scaled speciation times *T*_1_ = 1.6, *T*_2_ = 2.4 (scaled by *θ*_0_ = 0.005), coalescent rates *C*_1_ = *C*_2_ = *C*_3_ = *C*_4_ = 1 and gene flow rates *M*_1_ = *M*_2_ = 0.5. The true scaled speciation times *T*_1_ = 1.6, *T*_2_ = 2.4 were used to identify different gene tree histories. The MLEs are *Ĉ* = 0.99, and 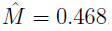, indicated by the white cross in the plot, with log likelihood *l* = −2001.745.

We repeated this simulation process 1,000 times for varying numbers of loci, and obtained the mean and standard deviation for the MLEs of C and M (Table 1, section labeled “Simualtion Study 1”, columns 2 and 3). We see that the estimates of *C* and *M* appear to be generally unbiased, with increasing variance as the number of loci decreases. We also consider the case in which we are reasonably confident that *C*_1_ = *C*_2_ = *C*_3_ = *C*_4_ = 1 and wish to estimate *M*_1_ and *M*_2_ separately (Table 1, section labeled ”Simualtion Study 1”, columns 4 and 5). Again, our results suggest very good performance of our method in estimating the rates of gene flow in a three-species model, with unbiased estimates and decreasing variance as the number of loci increases. Notably, in our simulation study, we only considered two parameters at a time in order to reduce the computational burden and to run more replications. In empirical data analyses described below, these parameters are estimated together.

**Table 1:**
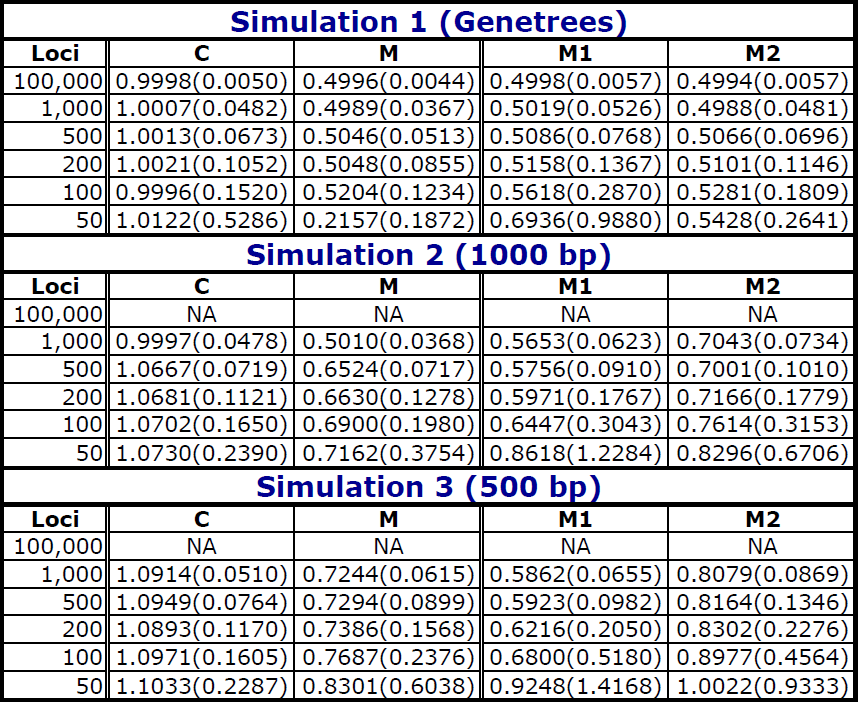
Maximum likelihood estimates of the scaled coalescent rates and the gene flow rates obtained from the simulated data sets under the three species coalescent model with gene flow between both sister taxa. The heading “Simulation 1” refers to the case in which gene trees are directly simulated from the given species tree by ms; “Simulation 2” refers to the case in which gene trees were estimated from 1,000bp sequences simulated by ms and then seq-gen; and “Simulation 3” refers to the case in which gene trees were estimated from 500bp sequences simulated by ms and then seq-gen. The columns labeled C and M refer to the MLEs of C and M under the assumption that *M*_1_ = *M*_2_ = *M*, and *C*_1_ = *C*_2_ = *C*_3_ = *C*_4_ = *C*. The columns labeled *M*_1_ and *M*_2_ refer to the MLEs of *M*_1_ and *M*_2_ when the scaled coalescent rates are fixed at their true values *C*_1_ = *C*_2_ = *C*_3_ = *C*_4_ = 1. All entries of the table are the mean over 1,000 repetitions of the simulation, with the standard deviation given in parentheses. The parameter values used to simulate the data in all cases were *C* = 1.0 and *M* = *M*_1_ = *M*_2_ = 0.5.

The first simulation study indicates good performance of our method when gene trees are known without error. However, in the typical empirical setting, gene trees must first be estimated from observed sequence data. Our second two simulation studies thus mimicked this condition by simulating sequence data and using gene trees estimated from these sequence data (using maximum likelihood in PAUP*) as the input into our method. We considered simulation data sets with either 1,000bp (Simulation Study 2) or 500bp (Simulation Study 3). The results of these simulations are shown in Table 1. It is reasonable that comparing with directly simulated gene trees, the gene trees estimated from the sequence data produce less accurate estimates for both parameters. However, when the number of loci is large enough (more than 200), our method can still produce good estimates using the DNA sequence data. Also, the simulations with sequences of lengths 1,000bp led to better estimates than the simulations with 500bp, since more information is provided with longer DNA sequences and the gene tree estimates should be more accurate for longer genes. An interesting finding is that the simulations using sequence data always overestimate the gene flow rates when we assume *M*_1_ *≠ M*_2_. More specifically, when we assume that *C*_1_ = *C*_2_ = *C*_3_ = *C*_4_ = 1, we notice that *M*_2_ is overestimated a lot, while *M*_1_ is only overestimated a little when there are more than 100 loci. However, when the directly simulated gene trees are used to estimate the gene flow rates, neither *M*_1_ and *M*_2_ is overestimated consistently. This finding indicates that information about ancient gene flow is more likely be lost during the process of estimating gene trees from sequence data.

### Application in an empirical Afrotropical mosquito data set

For the Afrotropical mosquito data set (Fontaine et al., 2015), we searched for the MLEs for *θ, M*_1_, and *M*_2_ in a two-step procedure. In the first step, we examined 60 equally-spaced values for *θ* ranging from 0.001 to 0.0594 and 60 equally-spaced values for both log(*M*_1_), and log(*M*_2_) ranging from -3 to 3, for a total of 360,000 values at which the likelihood was calculated. After finding that the likelihood was maximized along this grid at *M*_1_ = 0.1, *M*_2_ = 19.9526 and *θ* = 0.00675, we used a finer grid that consisted of 200 equally-spaced values of log(*M*_2_) ranging from -3 to 3, 200 equally-spaced values of *θ* ranging from 0.00396 to 0.01188, and four values of *M*_1_ (0.01, 0.1, 1, and 10). Figure 8 shows plots of the log likelihood for each level of *M*_1_, with red color indicating low log likelihood and blue color indicating high log likelihood. The MLEs found in this manner were 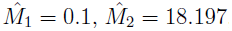 and 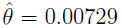. This result is highly consistent with the results of Fontaine et al. (2015), in which they conclude that there is substantial introgression between species *An. ara* and the ancestor of *An. col* and *An. gam* (Fontaine et al., 2015, Figure 1 (c)). The introgression between species *An. col* and *An. gam* is not tested in Fontaine et al. (2015), and our model suggested a lack of significant gene flow between these two species.

**Figure 8:**
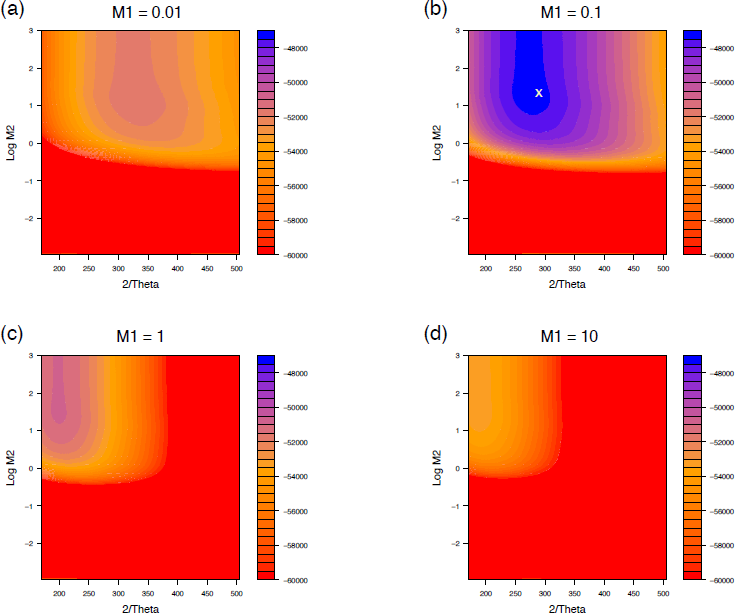
Contour plots of the log likelihood for the coalescent rate 2*/θ* and the gene flow rate *M*_2_ at four different levels of gene flow rate *M*_1_ assuming that *θ*_1_ = *θ*_2_ = *θ*_3_ = *θ*_4_ = *θ*. We analyzed 24,921 gene trees that were constructed from the whole genome alignment of the members of the *Anopheles gambiae* species complex. The estimated species tree topology and the speciation times were given by Fontaine et al. (2015). Four different levels of the gene flow rate between species A and B are shown in the panels: (a) *M*_1_ = 0.01; (b) *M*_1_ = 0.1; (c) *M*_1_ = 1; (d) *M*_1_ = 10. The MLEs are 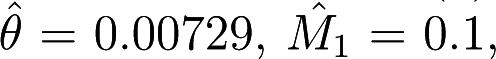, and 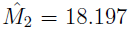, indicated by the white cross in panel (b), with log likelihood *l* = −47, 273.348.

This empirical study suggests that our model performs well in estimating the rates of coalescence and gene flow when the species tree, including speciation times, is well-estimated. We note that we used non-overlapping windows of 1000 kb as the “genes” for our study, and thus regions that were adjacent on the chromosome were used. This represents a violation of the basic model, in the sense that there may be recombination within our “genes” and a lack of recombination between “genes”. However, we feel that the use of such a large data set (nearly 25,000 genes) and the fact that recombination has been found to have a minor role in similar analyses that assume the absence of intralocus recombination (Lanier and Knowles, 2012) alleviates concern about this procedure. Our research validates the major introgression event between species *An. ara* and the ancestor of *An. col* and *An. gam*, in agreement with Fontaine et al. (2015). Our research also suggests that there is only a small amount of gene flow between species *An. col* and *An. gam*.

## DISCUSSION

### Identifying the species tree from the gene tree topology distribution in the presence of gene flow

In the coalescent model for three species with no gene flow following speciation, the gene tree topology that matches the species tree will always have probability that is at least as large as that of the other two topologies, with equality only occurring when the time between the two speciation events is 0. When gene flow occurs between sister species, however, there are portions of the parameter space in which all three gene tree topologies have equal probability, even when the time interval between speciation events is not 0 (see Figure 4(c) and Figure 5(l)). Interestingly, we can characterize the portions of the parameter space for which this happens by the following: when there is substantial gene flow between sister lineages deeper in the tree, then there must also be substantial gene flow between sister species near the tips of the tree in order for the species tree *topology* to be identifiable from the gene tree topology distribution (compare Figure 4(c) to Figures 4(f), (i), and (l); compare Figure 5(l) to Figures 5(c), (f), and (i)). This finding is sensible, because in the presence of a high rate of ancestral gene flow in the absence of gene flow elsewhere in the tree, the possible orders of coalescence among the three lineages will occur with approximately equal probability, and all three topologies will be equally likely, mimicking the case in which there is no gene flow and no time elapses between species events.

This finding has important implications for species tree estimation. Several new methods for estimating species trees from large data sets have used the “rooted triples” method to build trees for subsets of the overall data set, with a second step in which the trees based on triplets are assembled into an overall species tree estimate (Ewing et al., 2008; DeGiorgio and Degnan, 2010; Liu et al., 2010; Poormohammadi et al., 2014). This method is expected to work well in the absence of gene flow, because the rooted triple with the highest probability under the coalescent model is known to be displayed on the species tree (Degnan et al., 2009). However, our result shows that in presence of gene flow (and more specifically, in the presence of gene flow more ancestrally in the rooted triple but not between the sister taxa near the tips), the rooted triple relationships cannot be accurately inferred given only topology frequencies. Adding stochastic variance due to the mutation process could result in misidentification of the correct rooted triple, biasing species tree inference methods based on this assumption when gene flow is present.

Finally, we recall our earlier result concerning identifiability of coalescent parameters along a fixed species tree in the presence of gene flow, noting again the contrast with results in the absence of gene flow. In particular, in the absence of gene flow, the probability distribution of gene tree topologies identifies both the species tree topology and associated speciation times (Allman et al., 2011a). Here we showed that, even for a fixed species tree topology, many different coalescent parameter values may lead to the same distribution on gene tree topologies (Figure 6). This, too, has implications for species tree estimation, in that methods based only on the distribution of gene tree topologies cannot be used to infer species tree coalescent parameters. The distribution of gene tree histories, however, does appear to identify parameters, though we have not established this formally. This conjecture is supported by the positive performance of our method based on history distributions for both simulated and empirical data.

### Limitations in applying the method to empirical data

Though there are many benefits in using the distribution of gene tree histories to estimate the coalescent parameters and the rates of gene flow, the application of this method has some limitations for empirical data. First, in order to classify the gene tree genealogies into different gene tree histories, a species tree with known speciation times is required. Though the species tree could first be estimated from the data, this would greatly increase the computational cost and possibly lead to biases in the ultimate estimates if variability in the estimated species tree is not properly accounted for in the subsequent estimate of the coalescent parameters. However, when a good estimate of the species tree and corresponding speciation times is given (as in our example data set of Afrotropical mosquitos), our method can provide accurate estimates of the rates of gene flow and of the effective population sizes, parameters that are normally difficult to estimate.

Second, a fairly large number of loci are required to apply our method to an empirical data set. Our model provides the theoretical probability distribution of gene tree histories, but these probabilities are not directly observable in practice; rather, they must be estimated from the observed frequencies of gene tree histories estimated from data. As the number of loci increases, the distribution of the observed gene tree histories will be closer to the theoretical distribution, in the absence of error in estimating the gene tree genealogies. We showed the performance of our method in estimating parameters using different numbers of loci (Table 1). For simulated gene trees, at least 100 loci are required to give reasonable estimates of the parameters. For simulated DNA sequences, at least 200 loci are required. For empirical data, we suggest using as many loci as possible to get good estimates of the effective population sizes and the gene flow rates. In our example for the Afrotropical mosquito data set, 24,921 gene trees from 1 kb non-overlapping windows across the whole genome alignment were constructed, and the parameters estimated from this large data set were very reasonable and consistent with previous work (Fontaine et al., 2015).

Finally, there is a computational burden incurred when working with this model due to the need for matrix exponentiation and numerical integration. Computations are feasible when the values of the coalescent parameter *θ*_*x*_ and the speciation time *τ*_*y*_ are in a reasonable range. The reference species tree we used in this paper follows the parameters used in the research of Zhu and Yang (2012), with *θ*_*A*_ = *θ*_*B*_ = *θ*_*C*_ = *θ*_*AB*_ = 0.005, *τ*_1_ = 0.004, and *τ*_2_ = 0.006. We suggest keeping the ratio of *τ/θ* less than 10 to avoid any numerical issues. These issues can likely be overcome by implementing more sophisticated methods for calculating the matrix exponential and the numerical integrals.

### Future directions

Our model is constructed for three species with gene flow between both pairs of sister species. The model can be extended to an arbitrary number of species with gene flow between all sister-species pairs by constructing larger instantaneous rate matrices for each time period and increasing the dimension of integrals. In this case, there would also be more time intervals along the species tree that would need to be considered. For example, the largest instantaneous rate matrix for a four-species bifurcating tree will be 29 *×* 29, and it would require a three-dimensional integral. While it is not difficult to list all the density functions and the marginal probability functions as we have done here, the computational cost of calculating these quantities grows rapidly. A spectral decomposition method similar to that used in Andersen et al. (2014) could be an effective way to overcome the computational burden. In their paper, the spectral decomposition method was used to model a scenario in which an ancestral population splits into P subpopulations at some time *T*_*A*_ in the past (Andersen et al., 2014). A similar method of dividing a rate matrix into several submatrices could be helpful in implementing our model for an arbitrary number of species.

Another extension of our model is to add more sequences for each species. As in the case of adding more species discussed above, adding more lineages will also increase the dimension of the integrals as well as the size of the instantaneous rate matrices. Again, the main issue is improving the computational method so that the numerical probabilities of each gene tree history can be calculated efficiently.

A further extension of our model would be a model in which gene flow can occur globally throughout the phylogeny, rather than simply between sister species. This would increase biological realism, because though it is possible that most gene flow happens between closely related species, it is also possible that gene flow exists in more distantly-related species. As shown in Figure 1, our model assumes no gene flow between species A and C, and species B and C from the present to time *τ*_1_. If gene flow existed to some extent between species A and C, and species B and C, additional gene tree histories would be possible, and the symmetric probabilities of topologies G2 and G3 may be affected by the induced gene flow. To implement a model with more widespread gene flow, we need to introduce a new instantaneous rate matrix to describe the coalescent and the gene flow process in the time interval from the present to time *τ*_1_, and then carry out calculations along the lines of what we did here for that new matrix. While the methods are straightforward, computational challenges are the main limitation of this approach.

### Conclusions

This article presents a method for computing the gene tree history distribution under the coalescent model for three species that allows gene flow between both sister populations. The ability to compute gene tree history distributions for species trees with various effective population sizes as well as various gene flow rates leads to a better understanding of evolutionary relationships among closely related populations or species. The application of the gene tree history distributions in simulation studies as well as to empirical data sets allows us to infer species trees parameters, such as the ancient effective population sizes and the gene flow rates, using maximum likelihood. This study also demonstrates that for a fixed species tree topology, many different coalescent parameter values may lead to the same distribution on gene tree topologies, while the distribution of the gene tree histories is distinct for different choices of parameters. These findings have implications for the development of species tree estimation methods in the presence of gene flow. Future work is needed to formally establish conditions for identifiability of the species tree from the gene tree history distribution, as well as to extend the coalescent model with gene flow to an arbitrary number of species with more than one sampled genes per species.

## ACKNOWLEDGEMENT

We thank Drs. Matthew Hahn and James Pease for their willingness to share the mosquito data, and Dr. Pease for assistance in providing the data. Both authors were partially supported by NSF DEB-1455399 to L.K. and Dr. Andrea Wolfe.

## APPENDIX

For G1H4, the first coalescent event occurs after time *τ*_1_, while the second coalescent event occurs after time *τ*_2_, and thus *τ*_1_ *< t*_1_ *< τ*_2_ *< t*_2_. From the present to time *τ*_1_, no coalescent event occurs, and the probability is

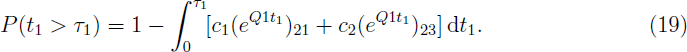

Similar to history G1H3, after time *τ*_1_, there are four distinct ways in which the two coalescent events can happen. In all four cases, the process starts in state ddc. The first case, denoted by G1H4C1, goes from state ddc to ddd, and then a coalescent event occurs and the state is dd. The final coalescent event does not occur before time *τ*_2_. Thus the state can change to any of the three states: dd, dc, and cc. To model this, we use Q3 to calculate the change from state ddc to ddd, and use Q2 for the change from state dd to dd, dc or cc. The density function is

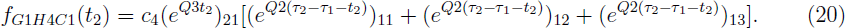

Similarly, we can write the density functions for the other three probabilities. The second probability has the sequence of states ddc - ddc - dc - dd/dc/cc, with corresponding density function

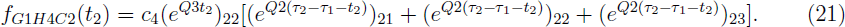

The third probability has the sequence of states ddc - ccc - cc - dd/dc/cc, with corresponding density function

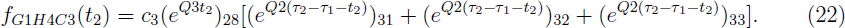

Finally, the fourth probability has the sequence of states ddc - ddc - dc - dd/dc/cc, with corresponding density function

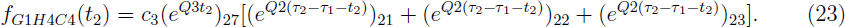

Thus the overall density function for *t*_1_ and *t*_2_ for history G1H4 is

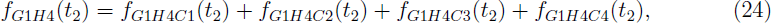

and the marginal probability for G1H4 is

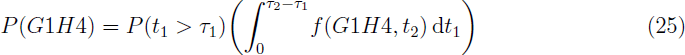

For G1H5, both coalescent events occur after time *τ*_2_ and the lineages that come from species A and B coalesce first, thus *τ*_2_ *< t*_1_ *< t*_2_. After time *τ*_2_, any two lineages have the same probability of coalescing. Since no coalescent events occur before time *τ*_1_, after time *τ*_1_ the state can change from ddc to all eight possible states. Thus, the probability of G1H5 is

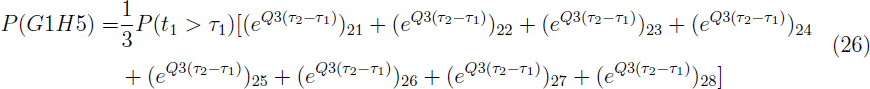

History G2H1 can be analyzed with a procedure similar to that used for history G1H3, and the probability of G2H1 is

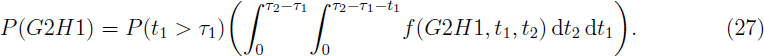

Here

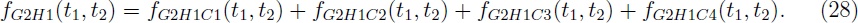

In the case of G2H1, some of the state changes are different than G1H3. The first case still goes from the state ddc to ddd, with corresponding density function

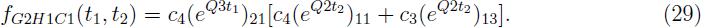

The second case has the sequence of states ddc - dcc - dc - dd/cc, with corresponding density function

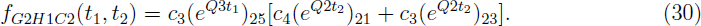

The third probability has the sequence of states ddc - ccc - cc - dd/cc, with corresponding density function

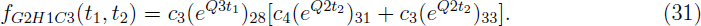

The last probability has the sequence of states ddc - dcc - dc - dd/cc, with corresponding density function

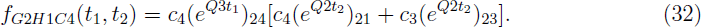

Similar to history H1G4, we can write the probability of history G2H2:

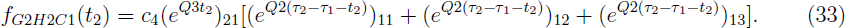

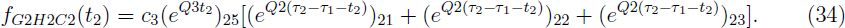

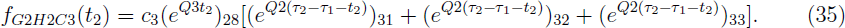

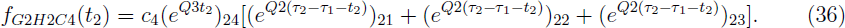

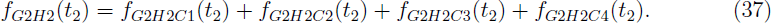

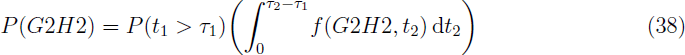

When both coalescent events occur after time *τ*_2_, any two lineages will have the same probability of coalescing, thus the probability of history G2H3 should be exactly the same as history G1H5. Also, due to the symmetry between G3H1 to G3H3 and G2H1 to G2H3, the probabilities of G3H1, G3H2, and G3H3 should be equal to the probabilities of G2H1, G2H2, and G2H3, respectively. Thus we have

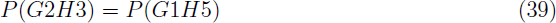

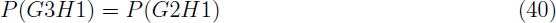

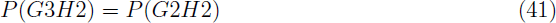

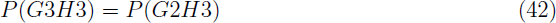

## Supplemental Figures

**Figure S1:**
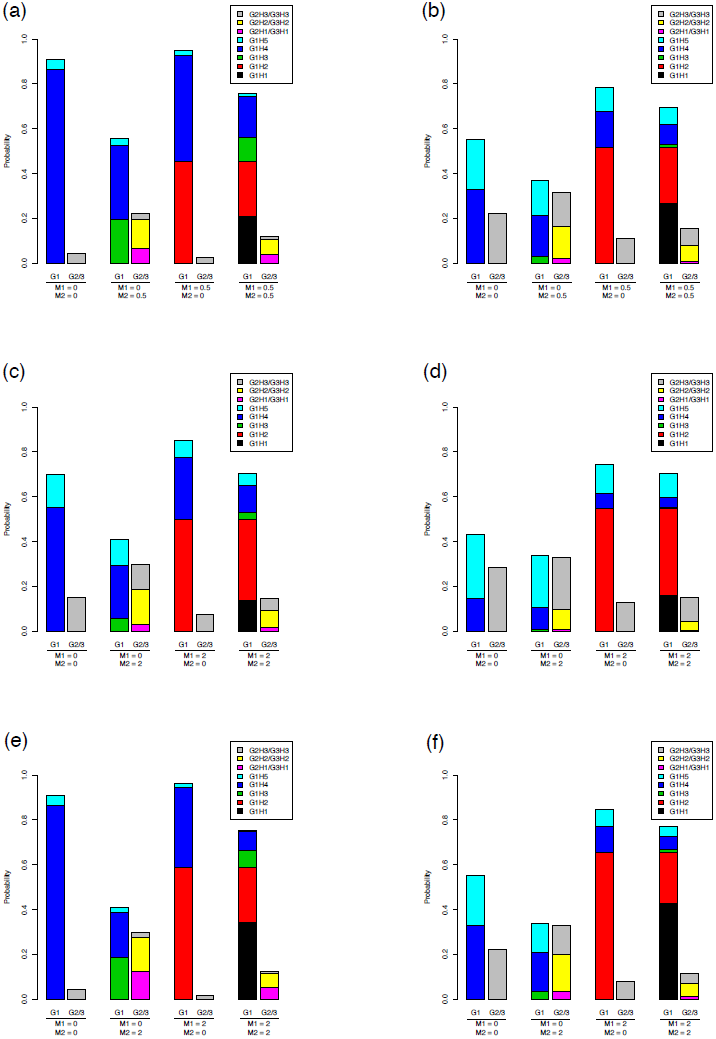
The probability distribution of the gene tree histories under species trees with scaled speciation times *T*_1_ = 1.6, *T*_2_ = 2.4 (*τ*_1_ = 0.004, *τ*_2_ = 0.006, *θ*_0_ = 0.005) for panels (c) and (d), and *T*_1_ = 2, *T*_2_ = 4 (*τ*_1_ = 0.01, *τ*_2_ = 0.02, *θ*_0_ = 0.01) for panels (a), (b), (e) and (f). Each gene tree history is denoted by a different color as shown in the figure. The probability of topology G1 is shown by the height of the column labeled G1; the height of the column labeled G2/3 shows the equal probability of the topologies G2 and G3. Thus, *P*(*G*1) + 2*P*(*G*2*/*3) = 1. The two sets of the scaled coalescent rates are *C*_1_ = *C*_2_ = *C*_3_ = *C*_4_ = 1 (scaled by *θ*_0_ = 0.005) in panels (a), (c), and (e), and *C*_1_ = 1, *C*_2_ = *C*_3_ = 0.5, *C*_4_ = 0.2 (scaled by *θ*_0_ = 0.005) in panels (b), (d) and (f). Each panel contains four cases of different gene flow rates: 1, no gene flow (*M*_1_ = *M*_2_ = 0); 2, no gene flow between species A and B (*M*_1_ = 0, *M*_2_ = 0.5 for panels (a) and (b); *M*_1_ = 0, *M*_2_ = 2 for panels (c) - (f)); 3, no gene flow between species C and the ancient species AB (*M*_1_ = 0.5, *M*_2_ = 0 for panels (a) and (b); *M*_1_ = 2, *M*_2_ = 0 for panels (c) - (f)); 4, equal rates of gene flow in both sister species (*M*_1_ = *M*_2_ = 0.5 for panels (a) and (b); *M*_1_ = *M*_2_ = 2 for panels (c) - (f)).

**Figure S2:**
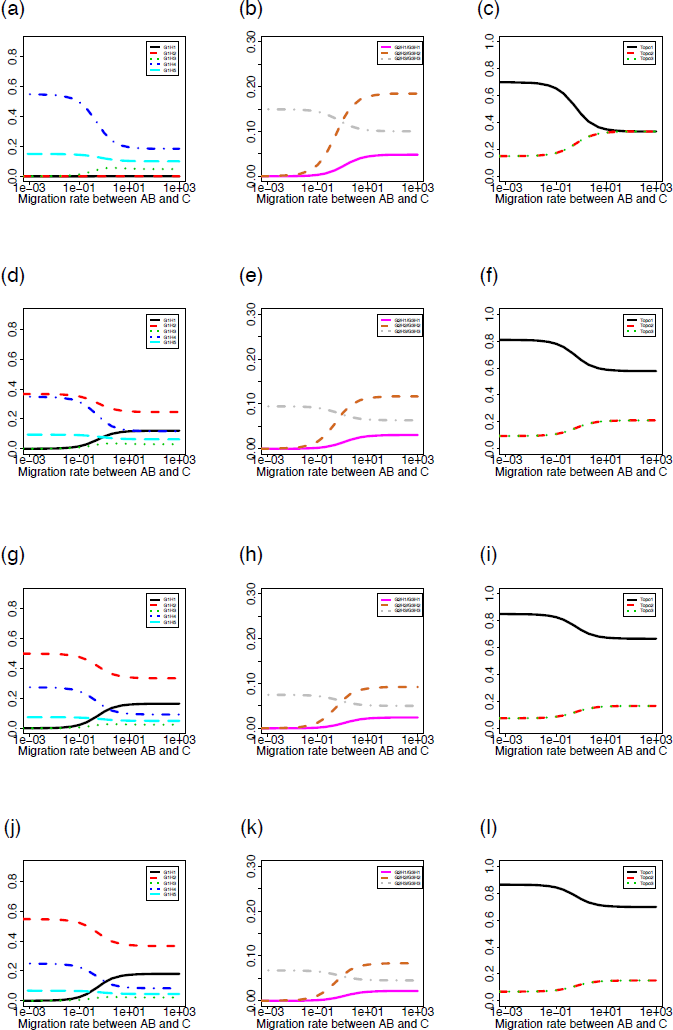
Probability distributions of the gene tree histories for the model of three species with gene flow between sister species. The probabilities of each gene tree history (y-axis) were plotted against the gene flow rate between species C and the ancient species AB *M*_2_ (x-axis; shown on a log scale). The four sets of parameter values are *C*_1_ = *C*_2_ = *C*_3_ = *C*_4_ = 1, *T*_1_ = 1.6, *T*_2_ = 2.4, *θ*_0_ = 0.005, *M*_1_ = 0.001 for panels (a) - (c); *C*_1_ = *C*_2_ = *C*_3_ = *C*_4_ = 1, *T*_1_ = 1.6, *T*_2_ = 2.4, *θ*_0_ = 0.005, *M*_1_ = 0.5 for panels (d) - (f); *C*_1_ = *C*_2_ = *C*_3_ = *C*_4_ = 1, *T*_1_ = 1.6, *T*_2_ = 2.4, *θ*_0_ = 0.01, *M*_1_ = 2 for panels (g) - (i); and *C*_1_ = *C*_2_ = *C*_3_ = *C*_4_ = 1, *T*_1_ = 1.6, *T*_2_ = 2.4, *θ*_0_ = 0.005, *M*_1_ = 20 for panels (j) - (l).

**Figure S3:**
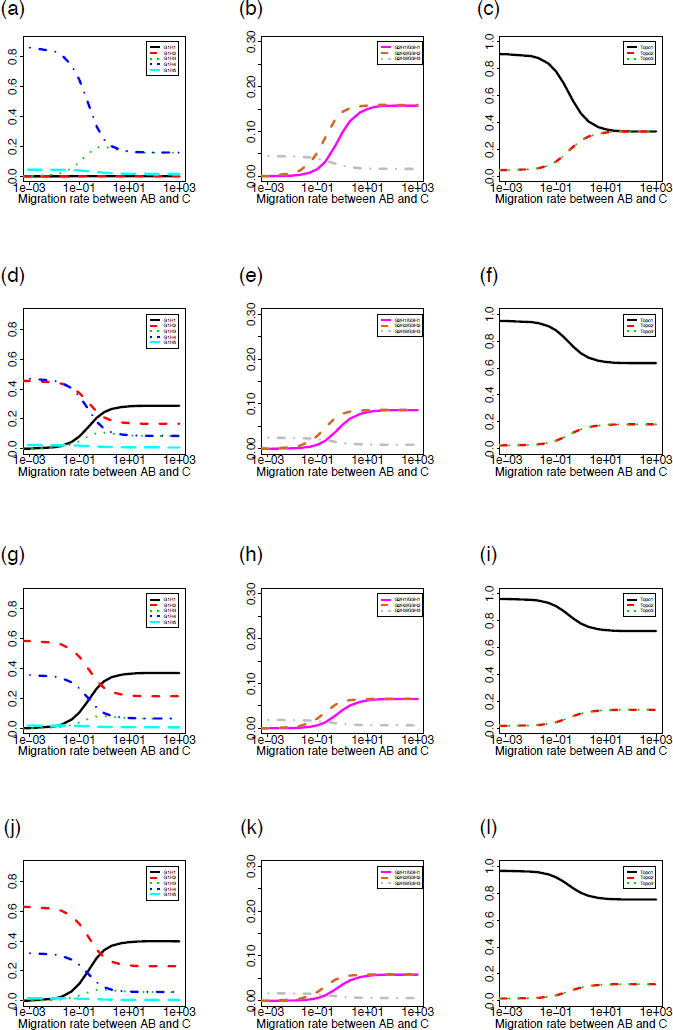
Probability distributions of the gene tree histories for the model of three species with gene flow between sister species. The probabilities of each gene tree history (y-axis) were plotted against the gene flow rate between species C and the ancient species AB *M*_2_ (x-axis; shown on a log scale). The four sets of parameter values are *C*_1_ = *C*_2_ = *C*_3_ = *C*_4_ = 1, *T*_1_ = 2, *T*_2_ = 4, *θ*_0_ = 0.005, *M*_1_ = 0.001 for panels (a) - (c); *C*_1_ = *C*_2_ = *C*_3_ = *C*_4_ = 1, *T*_1_ = 2, *T*_2_ = 4, *θ*_0_ = 0.005, *M*_1_ = 0.5 for panels (d) - (f); *C*_1_ = *C*_2_ = *C*_3_ = *C*_4_ = 1, *T*_1_ = 2, *T*_2_ = 4, *θ*_0_ = 0.01, *M*_1_ = 2 for panels (g) - (i); and *C*_1_ = *C*_2_ = *C*_3_ = *C*_4_ = 1, *T*_1_ = 2, *T*_2_ = 4, *θ*_0_ = 0.005, *M*_1_ = 20 for panels (j) - (l).

**Figure S4:**
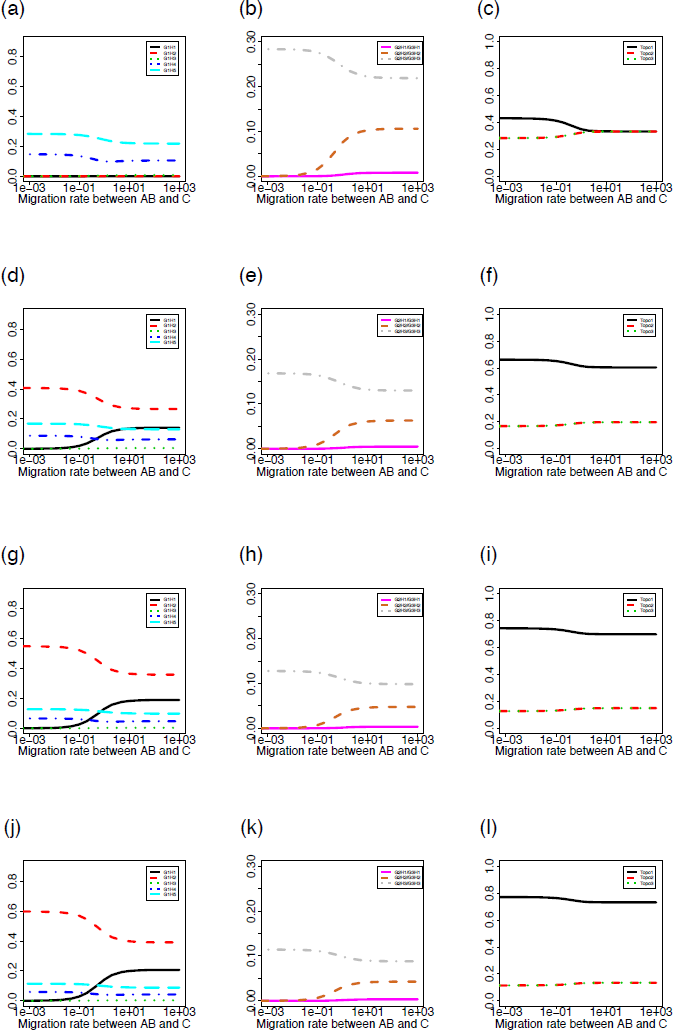
Probability distributions of the gene tree histories for the model of three species with gene flow between sister species. The probabilities of each gene tree history (y-axis) were plotted against the gene flow rate between species C and the ancient species AB *M*_2_ (x-axis; shown on a log scale). The four sets of parameter values are *C*_1_ = 1, *C*_2_ = *C*_3_ = 0.5, *C*_4_ = 0.2, *T*_1_ = 1.6, *T*_2_ = 2.4, *θ*_0_ = 0.005, *M*_1_ = 0.001 for panels (a) - (c); *C*_1_ = 1, *C*_2_ = *C*_3_ = 0.5, *C*_4_ = 0.2, *T*_1_ = 1.6, *T*_2_ = 2.4, *θ*_0_ = 0.005, *M*_1_ = 0.5 for panels (d) - (f); *C*_1_ = 1, *C*_2_ = *C*_3_ = 0.5, *C*_4_ = 0.2, *T*_1_ = 1.6, *T*_2_ = 2.4, *θ*_0_ = 0.01, *M*_1_ = 2 for panels (g) - (i); and *C*_1_ = 1, *C*_2_ = *C*_3_ = 0.5, *C*_4_ = 0.2, *T*_1_ = 1.6, *T*_2_ = 2.4, *θ*_0_ = 0.005, *M*_1_ = 20 for panels (j) - (l).

**Figure S5:**
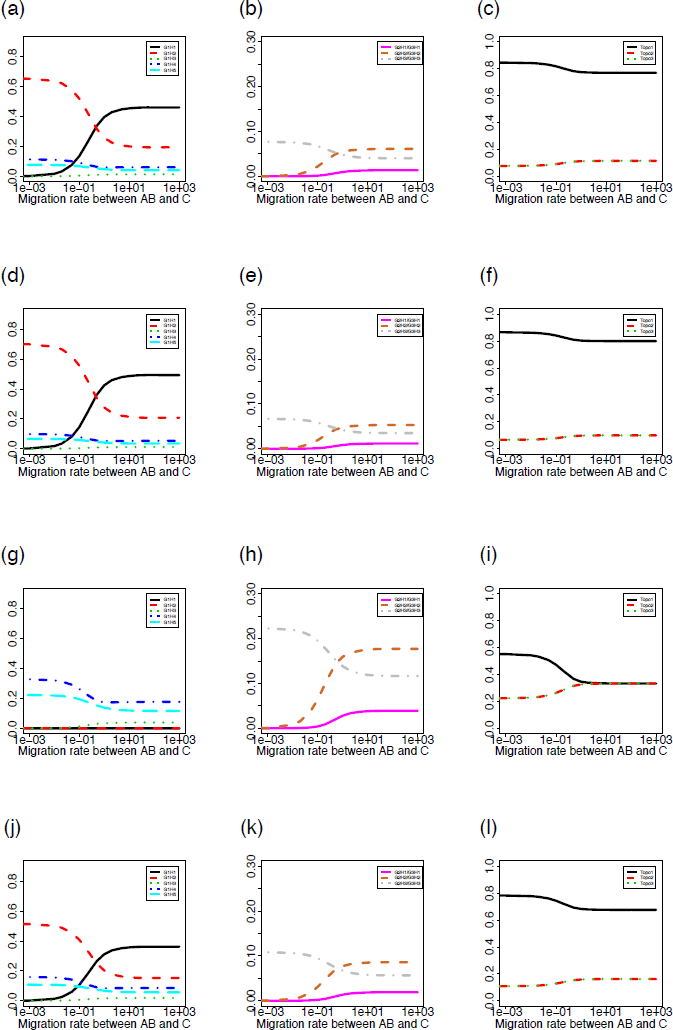
Probability distributions of the gene tree histories for the model of three species with gene flow between sister species. The probabilities of each gene tree history (y-axis) were plotted against the gene flow rate between species C and the ancient species AB *M*_2_ (x-axis; shown on a log scale). The four sets of parameter values are *C*_1_ = 1, *C*_2_ = *C*_3_ = 0.5, *C*_4_ = 0.2, *T*_1_ = 2, *T*_2_ = 4, *θ*_0_ = 0.005, *M*_1_ = 0.001 for panels (a) - (c); *C*_1_ = 1, *C*_2_ = *C*_3_ = 0.5, *C*_4_ = 0.2, *T*_1_ = 2, *T*_2_ = 4, *θ*_0_ = 0.005, *M*_1_ = 0.5 for panels (d) - (f); *C*_1_ = 1, *C*_2_ = *C*_3_ = 0.5, *C*_4_ = 0.2, *T*_1_ = 2, *T*_2_ = 4, *θ*_0_ = 0.01, *M*_1_ = 2 for panels (g) - (i); and *C*_1_ = 1, *C*_2_ = *C*_3_ = 0.5, *C*_4_ = 0.2, *T*_1_ = 2, *T*_2_ = 4, *θ*_0_ = 0.005, *M*_1_ = 20 for panels (j) - (l).

**Figure S6:**
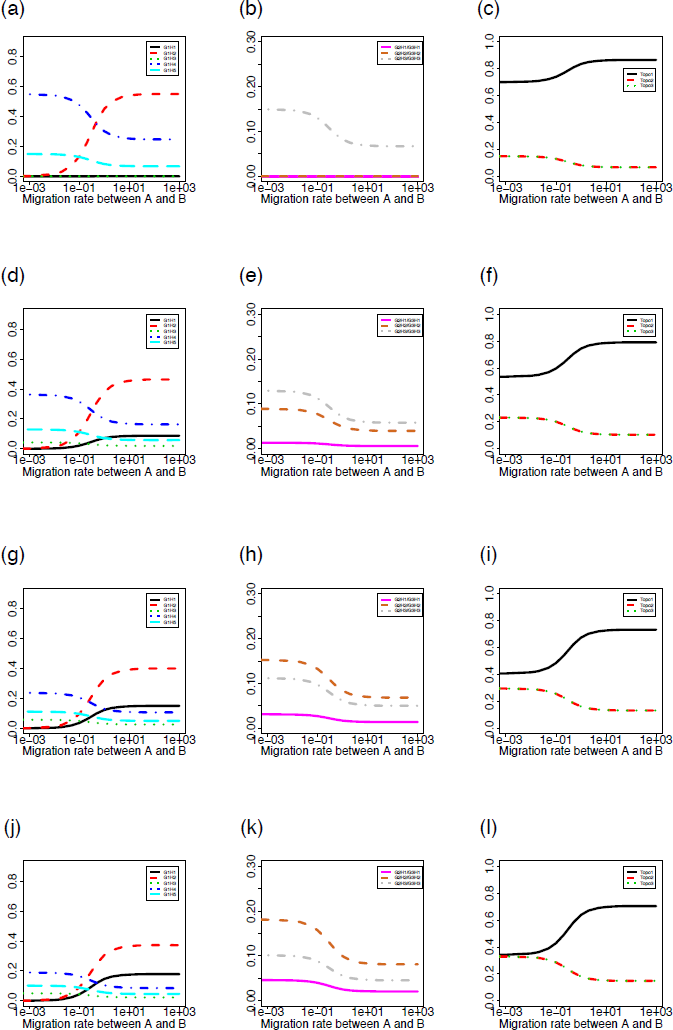
Probability distributions of the gene tree histories for the model of three species with gene flow between sister species. The probabilities of each gene tree history (y-axis) were plotted against the gene flow rate between species A and species B *M*_1_ (x-axis; shown on a log scale). The four sets of parameter values are *C*_1_ = *C*_2_ = *C*_3_ = *C*_4_ = 1, *T*_1_ = 1.6, *T*_2_ = 2.4, *θ*_0_ = 0.005, *M*_2_ = 0.001 for panels (a) - (c); *C*_1_ = *C*_2_ = *C*_3_ = *C*_4_ = 1, *T*_1_ = 1.6, *T*_2_ = 2.4, *θ*_0_ = 0.005, *M*_2_ = 0.5 for panels (d) - (f); *C*_1_ = *C*_2_ = *C*_3_ = *C*_4_ = 1, *T*_1_ = 1.6, *T*_2_ = 2.4, *θ*_0_ = 0.005, *M*_2_ = 2 for panels (g) - (i); and *C*_1_ = *C*_2_ = *C*_3_ = *C*_4_ = 1, *T*_1_ = 1.6, *T*_2_ = 2.4, *θ*_0_ = 0.005, *M*_2_ = 20 for panels (j) - (l).

**Figure S7:**
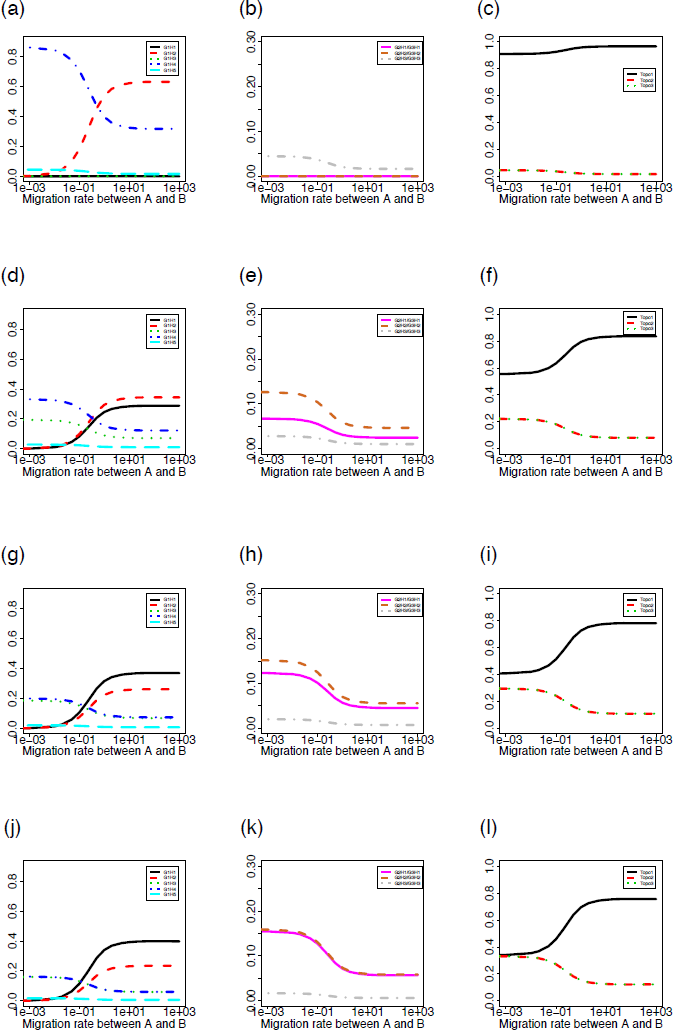
Probability distributions of the gene tree histories for the model of three species with gene flow between sister species. The probabilities of each gene tree history (y-axis) were plotted against the gene flow rate between species A and species B *M*_1_ (x-axis; shown on a log scale). The four sets of parameter values are *C*_1_ = *C*_2_ = *C*_3_ = *C*_4_ = 1, *T*_1_ = 2, *T*_2_ = 4, *θ*_0_ = 0.005, *M*_2_ = 0.001 for panels (a) - (c); *C*_1_ = *C*_2_ = *C*_3_ = *C*_4_ = 1, *T*_1_ = 2, *T*_2_ = 4, *θ*_0_ = 0.005, *M*_2_ = 0.5 for panels (d) - (f); *C*_1_ = *C*_2_ = *C*_3_ = *C*_4_ = 1, *T*_1_ = 2, *T*_2_ = 4, *θ*_0_ = 0.005, *M*_2_ = 2 for panels (g) - (i); and *C*_1_ = *C*_2_ = *C*_3_ = *C*_4_ = 1, *T*_1_ = 2, *T*_2_ = 4, *θ*_0_ = 0.005, *M*_2_ = 20 for panels (j) - (l).

**Figure S8:**
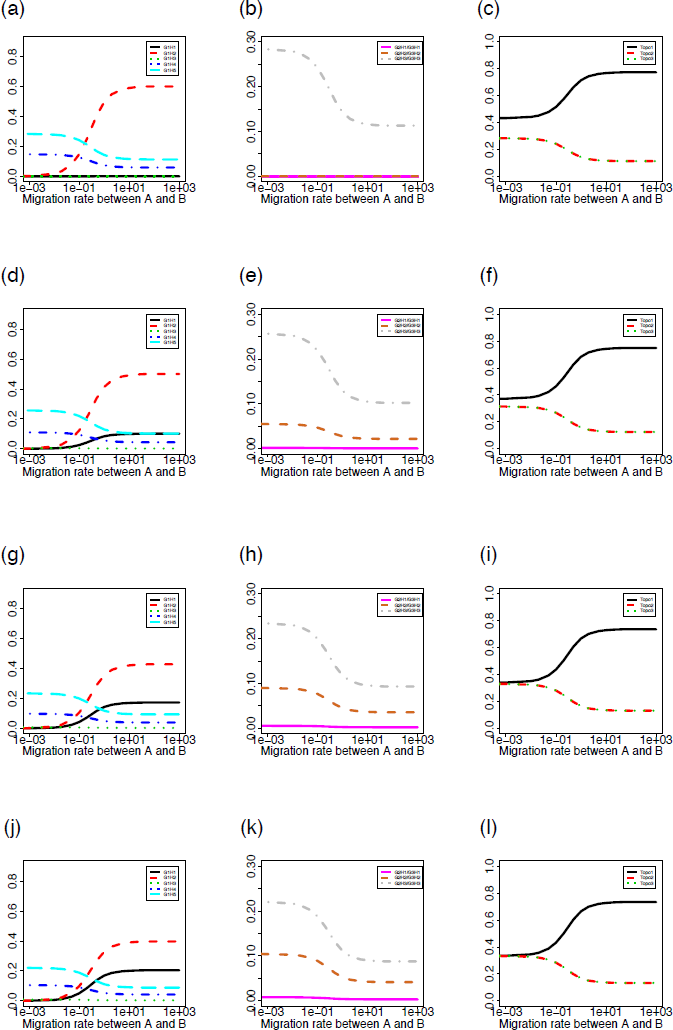
Probability distributions of the gene tree histories for the model of three species with gene flow between sister species. The probabilities of each gene tree history (y-axis) were plotted against the gene flow rate between species A and species B *M*_1_ (x-axis; shown on a log scale). The four sets of parameter values are *C*_1_ = 1, *C*_2_ = *C*_3_ = 0.5, *C*_4_ = 0.2, *T*_1_ = 1.6, *T*_2_ = 2.4, *θ*_0_ = 0.005, *M*_2_ = 0.001 for panels (a) - (c); *C*_1_ = 1, *C*_2_ = *C*_3_ = 0.5, *C*_4_ = 0.2, *T*_1_ = 1.6, *T*_2_ = 2.4, *θ*_0_ = 0.005, *M*_2_ = 0.5 for panels (d) - (f); *C*_1_ = 1, *C*_2_ = *C*_3_ = 0.5, *C*_4_ = 0.2, *T*_1_ = 1.6, *T*_2_ = 2.4, *θ*_0_ = 0.005, *M*_2_ = 2 for panels (g) - (i); and *C*_1_ = 1, *C*_2_ = *C*_3_ = 0.5, *C*_4_ = 0.2, *T*_1_ = 1.6, *T*_2_ = 2.4, *θ*_0_ = 0.005, *M*_2_ = 20 for panels (j) - (l).

**Figure S9:**
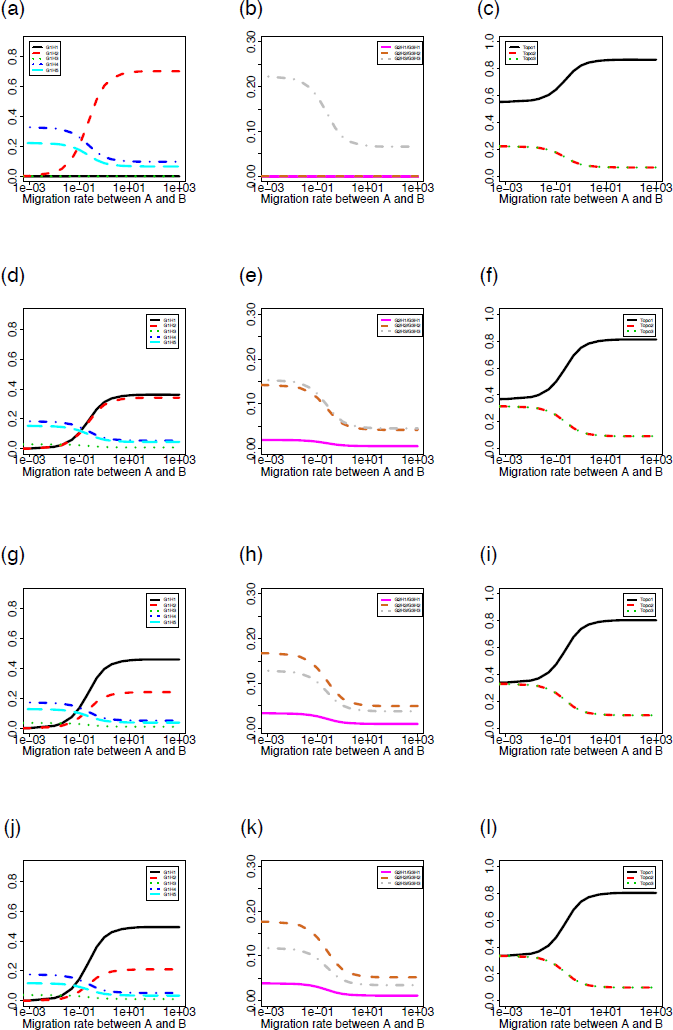
Probability distributions of the gene tree histories for the model of three species with gene flow between sister species. The probabilities of each gene tree history (y-axis) were plotted against the gene flow rate between species A and species B *M*_1_ (x-axis; shown on a log scale). The four sets of parameter values are *C*_1_ = 1, *C*_2_ = *C*_3_ = 0.5, *C*_4_ = 0.2, *T*_1_ = 2, *T*_2_ = 4, *θ*_0_ = 0.005, *M*_2_ = 0.001 for panels (a) - (c); *C*_1_ = 1, *C*_2_ = *C*_3_ = 0.5, *C*_4_ = 0.2, *T*_1_ = 2, *T*_2_ = 4, *θ*_0_ = 0.005, *M*_2_ = 0.5 for panels (d) - (f); *C*_1_ = 1, *C*_2_ = *C*_3_ = 0.5, *C*_4_ = 0.2, *T*_1_ = 2, *T*_2_ = 4, *θ*_0_ = 0.005, *M*_2_ = 2 for panels (g) - (i); and *C*_1_ = 1, *C*_2_ = *C*_3_ = 0.5, *C*_4_ = 0.2, *T*_1_ = 2, *T*_2_ = 4, *θ*_0_ = 0.005, *M*_2_ = 20 for panels (j) - (l).

